# Structural impact of 3-methylcytosine modification on the anticodon stem of a neuronally-enriched arginine tRNA

**DOI:** 10.1101/2024.11.18.624017

**Authors:** Kyle D. Berger, Anees M. K. Puthenpeedikakkal, David H. Mathews, Dragony Fu

## Abstract

All tRNAs undergo a series of chemical modifications to fold and function correctly. In mammals, the C32 nucleotide in the anticodon loop of tRNA-Arg-CCU and UCU is methylated to form 3-methylcytosine (m3C). Deficiency of m3C in arginine tRNAs has been linked to human neurodevelopmental disorders, indicating a critical biological role for m3C modification. However, the structural repercussions of m3C modification are not well understood. Here, we examine the structural effects of m3C32 modification on the anticodon stem loop (ASL) of human tRNA-Arg-UCU-4-1, a unique tRNA with enriched expression in the central nervous system. Optical melting experiments demonstrate that m3C modification can locally disrupt nearby base pairing within the ASL while simultaneously stabilizing the ASL electrostatically, resulting in little net change thermodynamically. The isoenergetic nature of the C32 – A38 pair vs the m3C32 – A38 pair may help discriminate against structures not adopting canonical C32 – A38 pairings, as most other m3C pairings are unfavorable. Furthermore, multidimensional NMR reveals that after m3C modification there are changes in hairpin loop structure and dynamics, the structure of A37, and the neighboring A31 – U39 base pair. However, these structural changes after modification are made while maintaining the shape of the C32 – A38 pairing, which is essential for efficient tRNA function in translation. These findings suggests that m3C32 modification could alter interactions of tRNA-Arg isodecoders with one or more binding partners while simultaneously maintaining the tRNA’s ability to function in translation.

## Introduction

Eukaryotic genomes encode hundreds of tRNA genes[1, 2]. Each tRNA can be grouped into an isoacceptor family based upon their anticodon sequence with all members in an isoacceptor family sharing the same anticodon sequence. The members within an isoacceptor family can be further divided into specific tRNA isodecoder species that each have the same anticodon, but contain sequence differences elsewhere in the tRNA body [3]. Even though each tRNA isodecoder within an isoacceptor family decodes the same codon, functional studies have identified differences in their biological activities implying that isodecoders do not simply serve redundant roles in translation [4, 5].

In mammals, the tRNA-Arg-UCU isoacceptor family contains 5 isodecoder members [1]. The tRNA-Arg-UCU-4-1 isodecoder (Figure 1A) is highly expressed in tissues encompassing the central nervous system (CNS) such as the brain and spinal cord, while exhibiting lower expression in organs outside the CNS [6]. The tRNA-Arg-UCU-4-1 isodecoder represents one of the most differentially expressed tRNA genes with >100-fold enrichment in the CNS compared to non-CNS tissues [7, 8]. In contrast, the 4 other tRNA-Arg-TCT isodecoders exhibit high expression in organs outside the CNS and low expression in the CNS. Notably, a point mutation in tRNA-Arg-UCU-4-1 causes decreased processing by RNaseP and reduced levels of mature tRNA-Arg-UCU [6, 9]. Reduction in tRNA-Arg-UCU-4-1 levels is associated with ribosome pausing at cognate AGA codons in the brain, activation of the integrated stress response, and neurodegeneration in mice [6, 10, 11].

**Figure 1.**
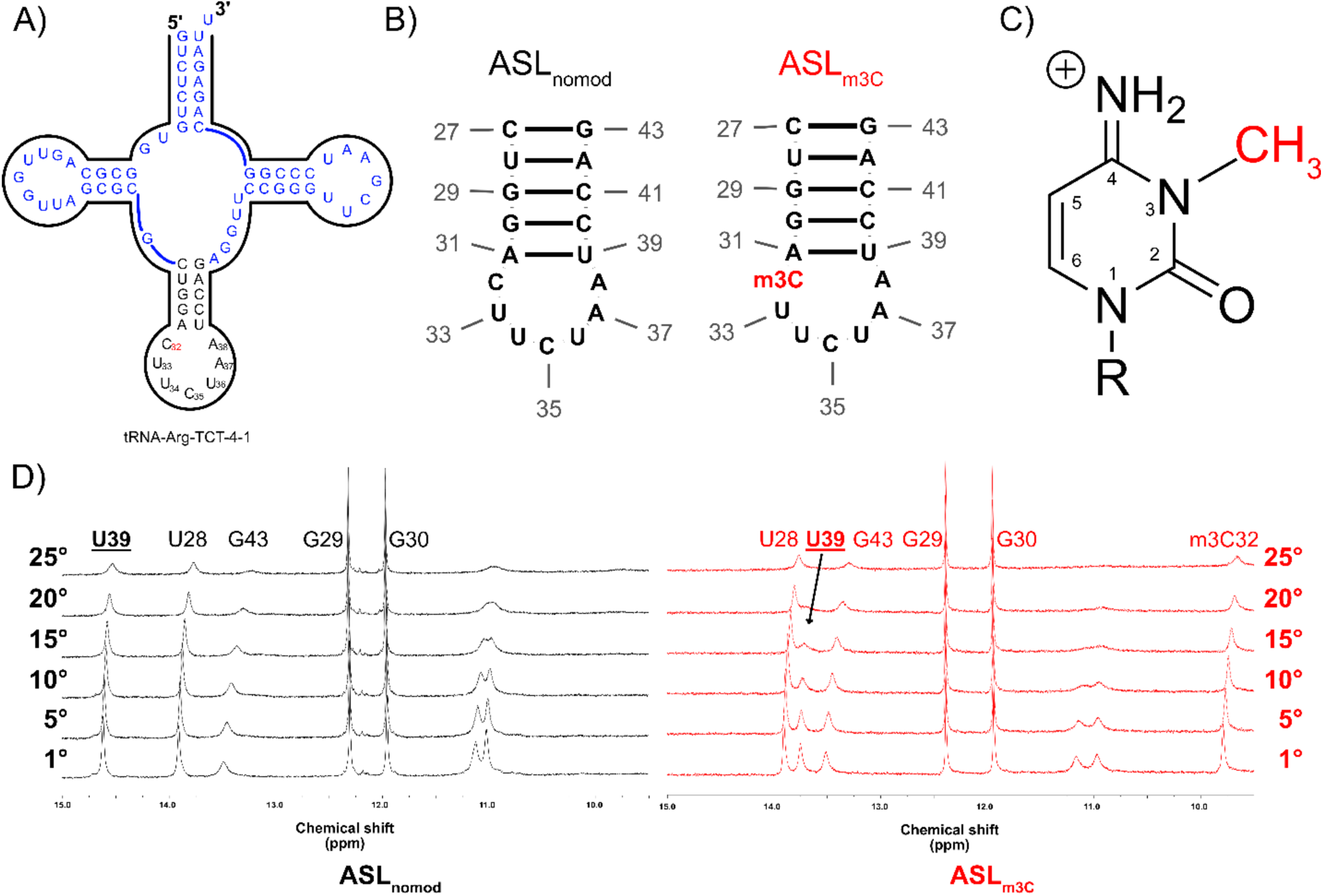
A: Sequence of neuronal tRNA-Arg-UCU-4-1 isodecoder displaying the anticodon stem loop in black. B: Anticodon stem loops ASLnomod and ASLm3C with their sequence and numbering scheme. C: Structure of the methylated m3C base with the numbering scheme used for referring to the m3C base. D: Imino proton spectra of ASLnomod and ASLm3C as a function of temperature. The peak labeled m3C32 results from the imine protons at position 4 within the m3C base.

The anticodon stem-loop (ASL) of mammalian tRNA-Arg-UCU isodecoders is subject to numerous chemical modifications that include 3-methylcytosine (m3C) at position 32, methoxycarbonylmethyl thiouridine (mcm5s2U) at position 34, and threonylcarbonyladenine (t6A) at position 37 [12–15]. The t6A and related 5-methylaminomethyluridine (mnm(5)U(34)) modifications are required for proper structure and binding to programmed ribosomes in the context of *E. coli* tRNA-Lys-UUU [16]. The t6A37 modification and other position 37 modifications such as hn6A are produced from a complicated biosynthetic pathway. Modification of A37 produces the open, structured loop required for ribosomal binding [17]. t6A modification at position 37 prevents U33-A37 base pair formation and alters positioning of U34 [17]. Both t6A and mcm5s2U modifications are required for pre-structuring, folding, and function of human tRNA-Lys-UUU [18]. The mcm5s2U modification increases stacking of nucleotides 34-35-36 of the anticodon, thus stabilizing the ubiquitous U-turn motif that is prevalent in ASLs [19, 20].

The m3C modification in tRNA-Arg-UCU is located in the anticodon loop at nucleotide position 32, which is known to form a noncanonical base-pair with nucleotide 38 in certain tRNAs [21–23]. The m3C methyl group resides on the N3 position of cytosine’s Watson-Crick pairing face. In unmodified cytosine-guanine pairs, this N3 position is involved in the central hydrogen bond with guanine and is flanked by a hydrogen bond on either side of N3. After m3C modification, all pairing schemes involving m3C are strongly destabilized, including the m3C-G pairing [24]. Accordingly, the m3C32 modification influences the structure of mt-tRNA-Ser-UCN and mt-tRNA-Thr ASLs both independently and acting in concert with (ms2)i6A37 and t6A37 modifications [25]. Notably, loss of m3C at position 32 in tRNA-Ser, Thr, and Arg of human and mouse cells causes alterations in gene expression and ribosome occupancy [26, 27]. These studies indicate that m3C modification on the ASL plays a key role in tRNA folding and decoding during translation [28].

Here, we present structures of the anticodon hairpin loop of CNS-enriched tRNA-Arg-UCU-4-1 in unmodified and m3C32 modified states. Our analyses reveal m3C32 modification in tRNA-Arg-UCU ASL has a surprising counterintuitive effect on RNA structure. It simultaneously stabilizes the ASL by relieving electrostatic repulsion while destabilizing local base pairing. NMR data suggests that m3C32 has a large effect upon structural dynamics within the ASL. Modification promotes dynamic anticodon behavior, destabilizing the nearby A31-U39 pair, structurally changes A37, and alters anticodon structure. Interestingly, however, the m3C32 – 38 pairing results in maintenance of the shape of the unmodified C32 – A38 canonical pairing. Thus, m3C modification results in an isosteric m3C32 – A38 pairing which allows the tRNA to maintain its shape at the 32-38 position while potentially fine-tuning other interactions through the ASL.

## Methods

### RNA Sample Preparation

Sequences of tRNA-Arg-UCU were retrieved from the tRNA database gtRNAdb [1, 2]. Thermodynamic analyses of the stability of the isolated ASL were performed with the aid of the RNA structure software [29] as well as manual calculation using the Nearest Neighbor Database (NNDB) [30]. Unmodified RNA (ASL_nomod_) was obtained from IDT with the sequence “5-CUGGA<u>CUUCUAA</u>UCCAG-3”. m3C modified RNA (ASL_m3C_) was obtained from Chemgenes with the sequence “5-CUGGA<u>(m3C)UUCUAA</u>UCCAG-3”. NMR samples were prepared by dissolution of the RNA within 500 μL of a buffer containing 10 mM sodium phosphate, 50 μM Na_2_ EDTA, and pH = 6.3. The final concentration of ASL_nomod_ and ASL_m3C_ within the RNA samples were 300 μM and 200 μM, respectively. Samples were annealed/folded by heating to 80°C and then slow cooling to room temperature upon the benchtop. D_2_O samples were prepared by lyophilization using a vacuum centrifuge and then subsequent re-dissolution in D_2_O. Lyophilization was repeated 3 times and on the final time, the sample was resuspended in 99.9% D_2_O to the original 500 μL.

### RNA Thermal Melting

Melting experiments were performed using a Shimadzu UV-1800 spectrophotometer. A slow temperature ramp of 1°C per minute between 20°C and 90°C was used with absorbance measured once per minute at a wavelength of 260 nm. To derive thermodynamic parameters comparable to the NNDB [30, 31], melting experiments were first performed in buffer containing 20 mM sodium phosphate, 1M NaCl, and pH = 7.0. Paradigms for extraction of hairpin thermodynamic parameters from thermal melting data have been described previously [32]. Thermal melting data was analyzed using the Meltwin and MeltR software [33, 34] assuming two-state melting. Generated thermodynamic data for the 7 nucleotide loop was compared to existing databases for thermodynamic data in 1M NaCl [30, 31]. In addition to the 1M NaCl buffer described previously, melting experiments were also undertaken in 3 separate buffers with NaCl concentration and pH conditions meant to mimic those in the cell. The buffers used a varying amount of magnesium while maintaining a constant pH of 7.5: (1: 20 mM sodium phosphate, 150 mM NaCl, 0 mM MgCl_2_. 2: 20 mM sodium phosphate, 150 mM NaCl, 1 mM MgCl_2_. 3: 20 mM sodium phosphate, 150 mM NaCl, 5 mM MgCl_2_). Lastly, a low pH buffer (pH = 5.0) composed of 150 mM NaCl and 20 mM MES was used to investigate the effect of pH on the stability of the ASL.

### NMR data collection

NMR data was collected on SUNY ESF’s 800 MHz Bruker spectrometer with data collection aid from SUNY ESF’s Analytical and Technical Services (ACTS). 1D spectra of the imino region were collected using a 3919 WATERGATE pulse sequence [35] for water suppression with a delay optimized for signals within the downfield imino region (10 – 15 ppm). 1H-1H NOESY spectra focused upon connection of imino protons to the remainder of the base/sugar region were also collected with 3919 WATERGATE water suppression. For both modified/unmodified RNAs, these imino focused NOESY spectra were collected at 5°C to maximize potential NOEs arising from imino protons. 1H-1H NOESY spectra focused on assignment of sequence backbone were collected with both a long (400 ms) and short mixing time (75 ms). Short mixing time (30 ms) 1H-1H WATERGATE TOCSY spectra were taken to aid in initial assignment of H5-H6 pyrimidine peaks. After samples were lyophilized and resuspended in D_2_O, higher resolution 1H-1H NOESY (short/long mixing times of 75/400 ms), 1H-1H TOCSY (30 ms), and natural abundance 13C-1H HSQC experiments were collected with minimal water suppression to aid in assignment of sugar pucker and sugar proton resonances.

### NMR analysis

Processing of spectra was accomplished using either Bruker’s TopSpin program and/or NMRpipe [36]. Processed data were converted into UCSF file format and analysis/assignment/peak integration proceeded within the NMRFAM-Sparky program [37].

### Structural modeling

Assigned NOESY spectra data were integrated using NMRFAM-Sparky. Integrated volumes were converted to distances using the relationship r=(constant/V)^6^. Restraints from peaks without overlap were calculated from integrated volumes by assuming 4-fold error within the NOE volume measurement. For overlapped peaks, restraints were determined based upon a binning approach where peaks were manually placed into bins based upon their apparent volume. Resonances were classified as strong (2.7 Å upper limit), medium (3.5 Å upper limit), weak (4.7 Å upper limit), or very weak by NOE volume (6 Å upper limit).

Dihedral restraints were assigned based upon typical values for A-form RNA, where applicable. The stem region (residues 27 – 31 and 39 – 43) was assumed to take an A-form conformation based upon the Watson-Crick pairing observed [38]. Dihedral restraints within the stem were given the following values: α = −65 ± 90°, β = 165 ± 75°, γ = 60 ± 60°, δ = 80 ± 35°, ε = −115 ± 125°, and ζ = −70 ± 90°. Residues within the loop that had a clear indication of C2’ endo preference (U34, C35, and U36) were restrained with δ = 130 ± 20°. Glycosidic angles were restrained to be in the anti-conformation with χ = 255 ± 85°. When applicable, lower bound restraints were assigned in the absence of a normal NOE.

Simulated annealing (SA) of both ASLs was accomplished with the OL3 force field within the Amber molecular dynamics package [39]. The starting structure for the unmodified ASL was generated using iFoldRNA v2 [40–42] and, for the modified ASL, ChimeraX [43, 44] was used. The system was subjected to a 5000-step energy minimization, in Generalized Born implicit solvent model [45] with a nonbonded interaction cutoff of 12 Å to resolve steric clashes. Then, 5 ps of dynamics were performed with an initial temperature of 3000 K and kept at 3000 K with a temperature coupling constant (TAUTP) of 0.4. In this step, the restraint weight was ramped from 0.1 to 1.0. The Berendsen temperature coupling algorithm [46] was used to keep the temperature constant. The temperature was then reduced to 100 K over 3 ps with TAUTP = 4.0, and then finally cooled to 0 K over 2 ps with TAUTP of 1.0 for 1 ps and varying linearly to 0.05 for the last 1 ps. The restraint weight was kept at 1.0 to enforce the NMR restraints throughout the cooling. The nonbonded cutoff was 15 Å and integration timestep of 1 fs was used throughout the dynamics. The penalty cutoff for NOE restraints was kept at −0.001 and salt concentration at 0.1M. A total of 2000 replicas were run and the 20 lowest energy structures with the zero torsion violations > 5 ° and zero distance violations > 0.1 Å.

### Force field parameter derivation for modified m3C base

For the m3C residue, the bond, angle, dihedral, improper rotation, and nonbonded parameters were assigned according to RNA.OL3 atom types. Charge parameters for m3C simulated annealing were determined using RESP [47] (Restrained Electrostatic Potential. For a more detailed description of RESP fitting, see supplemental methods section and leaprc/frcmod files). The Amber [39, 48] xleap GUI version was used to construct the m3C residue and Amber antechamber [49] was then used to generate a Gaussian input file for calculating the electrostatic potential. Gaussian [50] (version 16) HF 6-31-G* was used for geometry optimization and then to determine the electrostatic potential surrounding the m3C residue. Antechamber and Amber resp tools were used to fit the atom-centered point charges for the m3C base and C1’ atom. The remaining backbone atoms were included in the fit, but their charges were fixed at those corresponding to cytosine in the RNA OL3 forcefield [51]. These RESP charges were subsequently employed to construct frcmod file and lib file, which were used to make Amber input files, and simulated annealing was performed under distance and dihedral restraints. Fit charges for the m3C base are available within the supplemental methods section.

### Structure prediction

Predicted structures for ASL_nomod_ were determined using the recommended procedures for each respective prediction algorithm [52, 53]. If multiple predictions were outputted by the prediction algorithm, as is the case with most prediction algorithms, the structure with the lowest all-atom RMSD to ASL_nomod_ was chosen for comparison.

### Structural analysis

Molecular graphics and structural analyses utilized UCSF ChimeraX [43, 44]. In addition, analysis was also performed using Pymol [54] and cpptraj [55] within the amber suite. Drawing of secondary structures was accomplished with the aid of RNAcanvas [56]. Structures containing certain structural features (i.e. anti/syn adenine residues) were identified with RNA FRABASE [57]

## Results

### The effect of m3C modification on tRNA-Arg-UCU-4-1’s anticodon stem-loop stability

We used an NMR-based approach to investigate the structural effects of m3C modification on the anticodon-stem loop (ASL) of tRNA-Arg-UCU-4-1 (Figure 1A and B). We examined the ASL in the absence (ASL_nomod_) or presence of m3C modification at position 32 (ASL_m3C_). This approach has been used to determine the effect of other modifications on the ASL of tRNAs [17, 19, 58–62].

Thermodynamic predictions of ASL_nomod_ within tRNA-Arg-UCU-4-1 indicated that the hairpin loop was favorable in the absence of any modification (Figure S1), thus suggesting that ASL_nomod_ could independently fold into the desired hairpin structure [30, 31]. Hairpin loop thermodynamics are well estimated by the number of residues in the loop [63]. The identical modified sequence of ASL_m3C_ leads to the conclusion that hairpin loop formation in ASL_m3C_ would also be favorable.

Ability of the tRNA-Arg-UCU-4-1 ASL sequence to fold into a hairpin conformation was verified in two ways. First, UV melting data at 1M NaCl yielded thermodynamics independent of RNA concentration (Table S1), consistent with unimolecular folding. Melting in 1M NaCl revealed thermodynamics comparable to prediction for stability of the expected hairpin (Table 1) [30, 31]. The formation of the expected ASL hairpin conformation was further confirmed by dilution of the ASL_nomod_ NMR sample. Since duplex-hairpin equilibria can cause issues for self-complementary sequences [64, 65], formation of the expected ASL hairpin was further confirmed by NMR spectra of diluted samples. Spectra of diluted samples indicated a lack of concentration dependent change, further consistent with hairpin population within the ASL NMR samples (Figure S2).

**Table 1.** Melting characteristics of ASL_nomod_ and ASL_m3C_ at 1M NaCl.

| **Table 1. Melting characteristics of ASL<sub>nomod</sub> and ASL<sub>m3C</sub> at 1M NaCl** |  |  |  |  |
| --- | --- | --- | --- | --- |
| Construct | T<sub>m</sub> (°C) | $\Delta G^{\circ}_{37}$ (kcal/mol) | $\Delta H^{\circ}$ (kcal/mol) | $\Delta S^{\circ}$ (eu) |
| ASL-Arg-UCU<sub>nomod</sub> | 70.5±0.8 | -4.6±0.1 | -48.7±1.7 | -142±5 |
| ASL-Arg-UCU<sub>m3C</sub> | 69.3±0.5 | -4.0±0.2 | -42.3±1.4 | -132±4 |
| ASL-Arg-UCU<sub>pred</sub> |  | -4.05 |  |  |
| Note: Measurements performed in 1M NaCl. Derived values represent averages taken from individual fitting of at least 4 different RNA concentrations to a two-state model. Predicted values are obtained from the 2004 set of nearest neighbor parameters . |  |  |  |  |

Melting profiles of ASL_nomod_ and ASL_m3C_ were very similar at 1M NaCl (Table 1 and Figure S3). Even though the m3C modification disfavors canonical base pairing [24], lack of large changes in melting behavior is expected if C32’s pairing face is not in a position where insertion of a bulky methyl group would interfere with structure. Based upon the determined values for ΔG°_37_ at 1M NaCl (Table 1) and the expected secondary structure (Figure 1B), the sum of ΔG°_37_ (ΔG°_37_ (HP+TMM)) for the seven-nucleotide (nt) hairpin loop initiation (HP) and CA terminal mismatch (TMM) was calculated for ASL_nomod_ and ASL_m3C_ (Figure S4), yielding ΔG°_37_(HP+TMM) of 4.9 kcal/mol and 5.5 kcal/mol, respectively. This implies that the 7-nt hairpin loop and C32-A38 terminal mismatch is slightly more stable in ASL_nomod_ than in ASL_m3C_ at 1M NaCl. Comparison to the predicted value for ΔG°_37_(HP+TM) of ASL_nomod_, 5.4 kcal/mol), implies that the unmodified ASL_nomod_ hairpin loop/terminal mismatch is slightly more stable than predicted [30].

Melting experiments were also performed in 150 mM NaCl, varying Mg^2+^ concentration, and pH 7.5 to assess the effect of m3C modification on loop stability under conditions that more closely resemble the chemical environment within the cell. Under these conditions, ASL_nomod_ and ASL_m3C_ behaved differently from each other (Table 2). In contrast to 1M NaCl (without Mg^2+^), ASL_m3C_ becomes more stable than ASL_nomod_ at 150 mM NaCl (without Mg^2+^). When 1 or 5 mM Mg^2+^ is added in the presence of 150 mM NaCl, ASL_nomod_ and ASL_m3C_ stabilities are essentially identical within uncertainty (Table 2). One explanation for this behavior is easing of phosphate backbone electrostatic repulsion by the positively charged m3C32 base. Given that modification of cytosine into m3C within duplex regions is strongly destabilizing [24] but has little effect on stability at higher salt concentrations within ASL_m3C_ (Table 1 and Table 2), it is likely that m3C modification has differing effects depending on whether the cytosine being modified is in a pre-existing CG pair or already exists in a loop region.

**Table 2.**
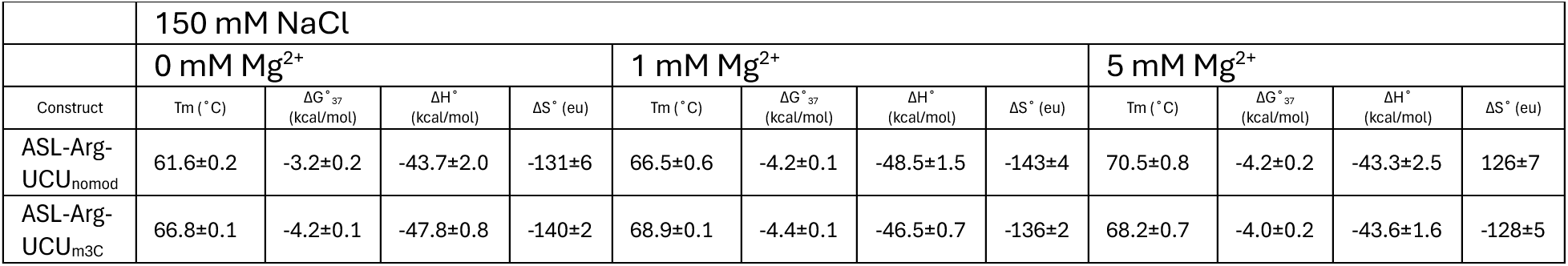
Melting characteristics of ASL_nomod_ and ASL_m3C_ at different Mg2+ concentrations.

A38 in the very similar ASL of tRNA-Lys-UUU has a pKa shifted to approximately 6.0 [58], resulting in a C_32_ − A^+^_38_ pairing (Figure S5). Sequence similarity between tRNA-Arg-UCU-4-1 and tRNA-Lys-UUU within the ASL led to the hypothesis that tRNA-Arg-UCU would also have a shifted A38 pKa. Furthermore, the effect of m3C32 modification across from the “paired” A38 (Figure 1B) could have large effects upon a potential C_32_ − A^+^_38_ pairing, as protonated CA pairs hydrogen bond through cytosine’s N3, which becomes blocked by the newly introduced methyl group in m3C (Figure 1C and S5). Melting experiments revealed the likely presence of a C_32_ − A^+^_38_ pairing in ASL_nomod_ by demonstrating a stabilization at lower pH (Table 3). ASL_nomod_ was stabilized at pH = 5.0, relative to pH = 7.5, as indicated by a shift in T_M_ from 61.6±0.2 °C at pH = 7.5 to 67.2±0.2 °C at pH = 5.0. A potential C_32_ − A^+^_38_ pairing could shift the structure of the ASL by facilitating an additional U33 – A37 pairing, resulting in formation of a hairpin triloop as is the case with tRNA-Lys-UUU [58]. Interestingly, acidification had the opposite effect on the T_M_ of ASL_m3C_. At pH = 7.5, the T_M_ of ASL_m3C_ is 66.8±0.1 °C, whereas at pH = 5.0 the T_M_ fell to a value of 59.5±0.3 °C (Table 3). This destabilization of ASL_m3C_ at low pH could be due to protonation of A38 like in ASL_nomod_, as the sequence surrounding position 38 (i.e. A37 and U39) remains the same. Since m3C introduces a positive charge, A38 protonation would lead to two positively charged bases directly across from one another adjacent to the stem. After modification, protonation at A38 in the presence of m3C32 would likely be disfavored due to a positive charge already existing on m3C32 and the m3C methyl group preventing similar pairing as C_32_ − A^+^_38_, thus leading to an acidic shift in the A38 pKa.

**Table 3.** Melting characteristics of ASL_nomod_ and ASL_m3C_ at low pH.

| Table 3. Melting characteristics of ASL<sub>nomod</sub> and ASL<sub>m3C</sub> at low pH |  |  |  |  |  |  |  |  |
| --- | --- | --- | --- | --- | --- | --- | --- | --- |
|  | pH 7.5 |  |  |  | pH 5.0 |  |  |  |
| Construct | T<sub>m</sub> (°C) | $\Delta G^{\circ}_{37}$<br>(kcal/mol) | $\Delta H^{\circ}$<br>(kcal/mol) | $\Delta S^{\circ}$<br>(eu) | T<sub>m</sub> (°C) | $\Delta G^{\circ}_{37}$<br>(kcal/mol) | $\Delta H^{\circ}$<br>(kcal/mol) | $\Delta S^{\circ}$ (eu) |
| ASL-Arg-UCU<sub>nomod</sub> | 61.6±0.2 | -3.2±0.2 | -43.7±2.0 | -131±6 | 67.2±0.2 | -3.7±0.3 | -41.6±3.5 | -122±10 |
| ASL-Arg-UCU<sub>m3C</sub> | 66.8±0.1 | -4.2±0.1 | -47.8±0.8 | -140±2 | 59.5±0.3 | -2.7±0.1 | -40.3±2.0 | -121±6 |

### Secondary structure of the tRNA-Arg-UCU-4-1 ASL_nomod_ and ASL_m3C_

Secondary structure of the ASLs was confirmed by 1D imino proton spectra (Figure 1D) and 2D NOESY data. The stem region showed evidence for all predicted base pairs (Figure 1D). Both ASL_nomod_ and ASL_m3C_ displayed strong resonances for all internal stem base pairs (U28-A42, G29-C41, G30-C40). The C27-G43 terminal pair’s imino proton resonance was assigned by process of elimination and its chemical shift corresponding with the expected value for a G imino in a GC pair [66]. Interestingly, the A31-U39 base pair (terminal base pair prior to start of the loop) displayed differential behavior depending on whether C32 was modified or not (Figure 1D). The U39 imino proton in ASL_m3C_ displays both a different temperature sensitivity and a change in chemical shift. The change in chemical shift is not unprecedented due to the different stacking environment caused by m3C modification [24]. However, the U39 imino resonance also displays less thermal stability in the presence of m3C32 (Figure 1D). This suggests that m3C32 modification induces more dynamics for U39. Destabilization of the canonical A31 – U39 pairing is interesting, as it demonstrates m3C not only exerts its effects upon its “paired” A38 residue, but also extends its effects into the stability of the adjacent stem region likely through differential stacking. Given that the ASL melting temperature and thermodynamics are similar before and after modification at higher salt concentrations (Table 1 and 2), there is likely some electrostatic relaxation provided by the positive charge of m3C32 at lower salt concentrations, thus offsetting destabilization of the A31 – U39 base pair induced by m3C32 in ASL_m3C_.

The ASL_nomod_ and ASL_m3C_ 1D imino proton resonances indicate significant dynamics for U33, U34, and/or U36 (Figure 1D). U33, U34, and/or U36 yield a cluster of broad proton resonances around 11 ppm (Figure 1D). These resonances could not be unambiguously identified and likely experience significant dynamics. NOE cross peaks arising from these resonances were not observed, indicating their exchange rate is too fast for observation of NOEs (as is often the case for unpaired iminos). Additionally, the imine proton resonance arising from position 4 on the m3C32 base was identified (Figure 1C and m3C32 labeled peak in 1D) by chemical shift and intra-residue NOEs to the methyl (m3C32 H4 – m3C32 H3) and H5 base proton (m3C32 H4 – m3C32 H5). These were the only NOEs observed from H4 of the m3C base, indicating that the H4 proton is unlikely to be involved in pairing.

To identify positions affected by m3C32 methylation, chemical shift perturbations were measured for ASL_m3C_ (Figure 2). Greater perturbations were observed in the loop and A31 – U39 closing base pair than in the rest of the stem. These perturbations suggest m3C modification triggers a structural rearrangement within the loop extending out to the anticodon itself. This implies non-nearest neighbor effects within the loop may affect anticodon structure and dynamics. For example, C35, an anticodon residue not positioned adjacent to the modified residue C32, displayed some of the largest observed shift changes (Figure 2).

**Figure 2.**
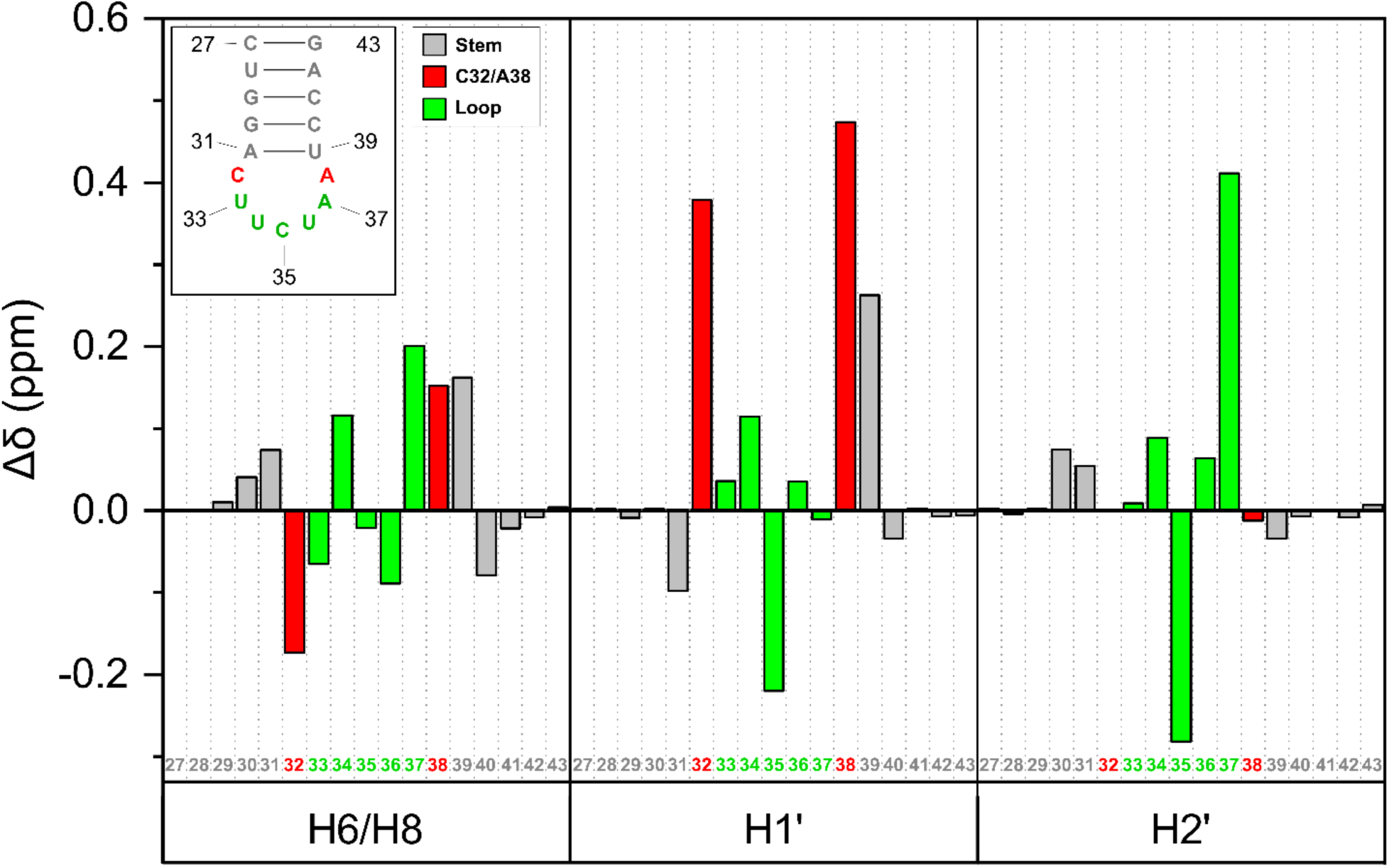
The change in chemical shift is shown on the Y-axis and the residue number and atom corresponding to that particular chemical shift are shown on the X-axis. Bars are colored according to whether they are part of the stem (white), terminal mismatch (red), or hairpin loop (green).

### A37 is strongly affected by m3C modification

Some of the larger structural changes observed center around A37 (Figures 2 and 3). In ASL_nomod_, A37 adopts a position that results in A37’s base coming into close contact with A38’s ribose H1’. This yields an unusually large inter-residue cross peak for A37 H1’ - A38 H8 (Figure 3A). In normal A-form helices, the distance between an n Ade H8 and n+1 Ade H1’ is 4.9±0.3 Å (Table S2) and the peak will only appear due to spin diffusion through the H2’. However, the NOE volume distance for A37 H1’ – A38 H8 was calculated as 2.9±0.3 Å in ASL_nomod_, indicating that the A37 – A38 connection is deviating from normal A-form. Interestingly, the A37 H2’ – A38 H8 peak was not similarly large relative to other n H2’ – n+1 H6/8 peaks nor was A37 H2 – A38 H1’. Modeling indicated that this A37 H1’ - A38 H8 restraint is satisfied by pinching of A38 base and A37 sugar together, resulting in significant tilt (Figure 3B). This differs from ASL_m3C_, where the A37 H1’ – A38 H8 distance is 3.6±0.4 Å, showing A37 in ASL_m3C_ still deviates from A-form, but not as greatly.

**Figure 3.**
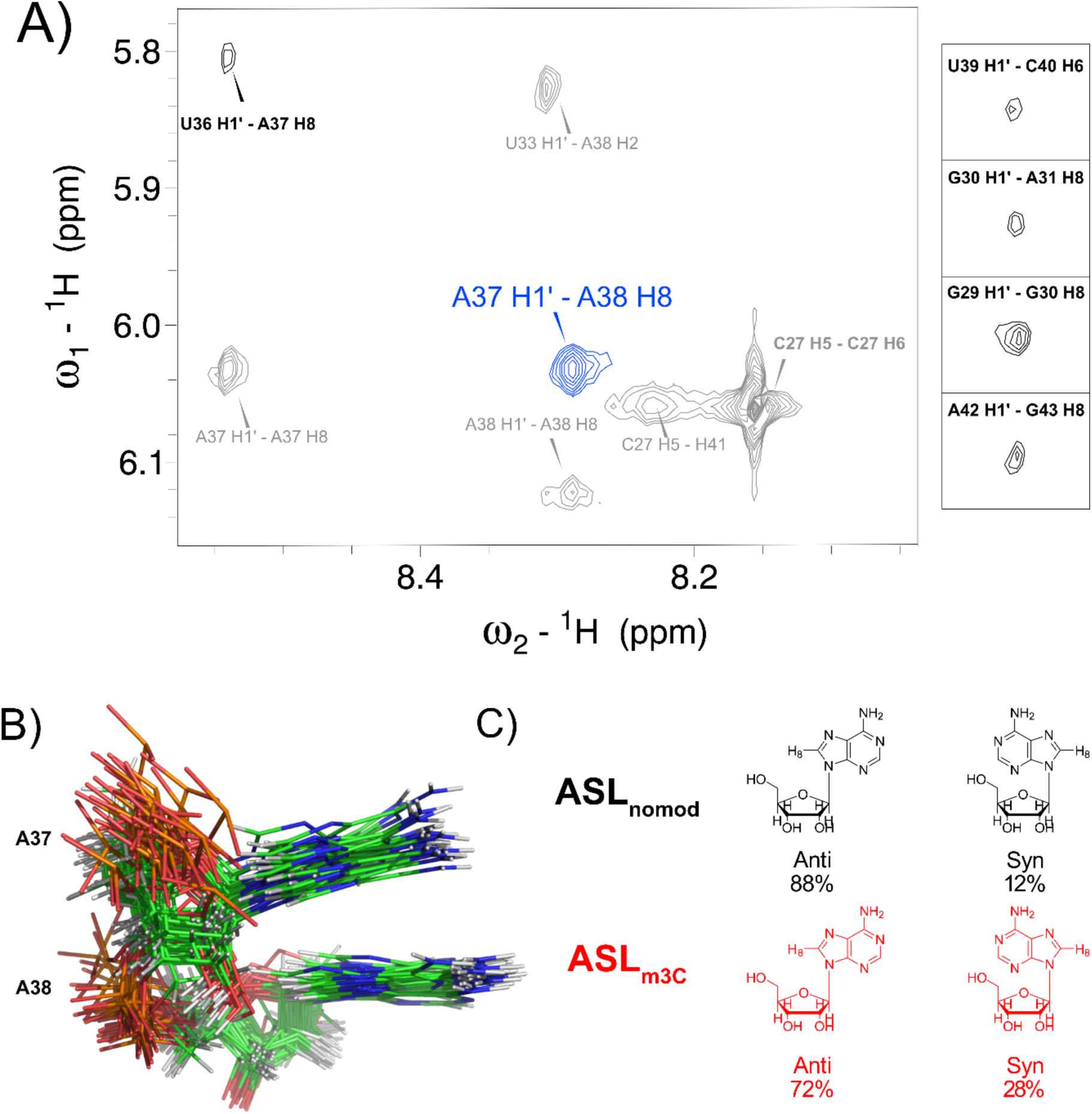
A: 75 ms NOESY spectrum of ASL_nomod_ displaying the unusually strong A37H1’ – A38 H8 NOE (blue). For comparison, other n H1’ – n+1 H8 NOEs (black) are shown to exemplify the average volume of similar resonances. B: Short A37 H1’ – A38 distance results in tilt between A37 and A38 residues. C: Structure of adenine bases in syn and anti conformations are shown along with the corresponding percentage of syn vs anti for ASL_nomod_ and ASL_m3C_.

A37 in ASL_nomod_ displays several NOEs to C35 that are not observed in ASL_m3C_. A37 H2 yields NOEs at short mixing time to C35 H5’ and C35H5’’, as well as weaker peaks to C35 H4’ and C35 H2’ (Figure S6). Furthermore, there is a A37 H1’ – C35 H5’ NOE that exists in ASL_nomod_. However, after modification in ASL_m3C_, this set of A37 – C35 peaks completely disappears. This indicates that m3C32 modification strongly influences A37, deterring long range interactions.

Both ASL_nomod_ and ASL_m3C_ have altered chi (χ) torsions as evidenced by shortened A37 H1’ – A37 H8 distances. A37 in ASL_nomod_ and ASL_m3C_ yield A37 H1’ – A37 H8 NOE distances of 3.3±0.3 Å and 3.0 Å, respectively. Given that pure syn and pure anti n H1’ – n H8 distances result in respective distances of 2.5±0.1 Å and 3.8±0.1 Å (Table S3), one would expect that A37 would on average occupy an intermediate chi torsion. Based upon pure syn and pure anti distances, the percentage of A37 adopting syn is 12% and 28% in ASL_nomod_ and ASL_m3C_, respectively (Figure 3C). The shorter n H1’ – n H8 A37 distance in ASLm3C would suggest that after modification the structure is pushed towards a more syn like conformation.

Further evidence of a structural transition for A37 is provided by a shift towards C3’ endo after modification (Figure 4A). While most other residues shifted towards a greater adoption of C2’ endo after modification, A37 was unique in that it shifted towards a C3’ endo sugar conformation after modification (Table 4 and Figure 4B). This shift towards C3’ endo may reflect the greater adoption of syn like character, as syn residues are less likely to adopt C2’ endo ribose [67].

**Figure 4.**
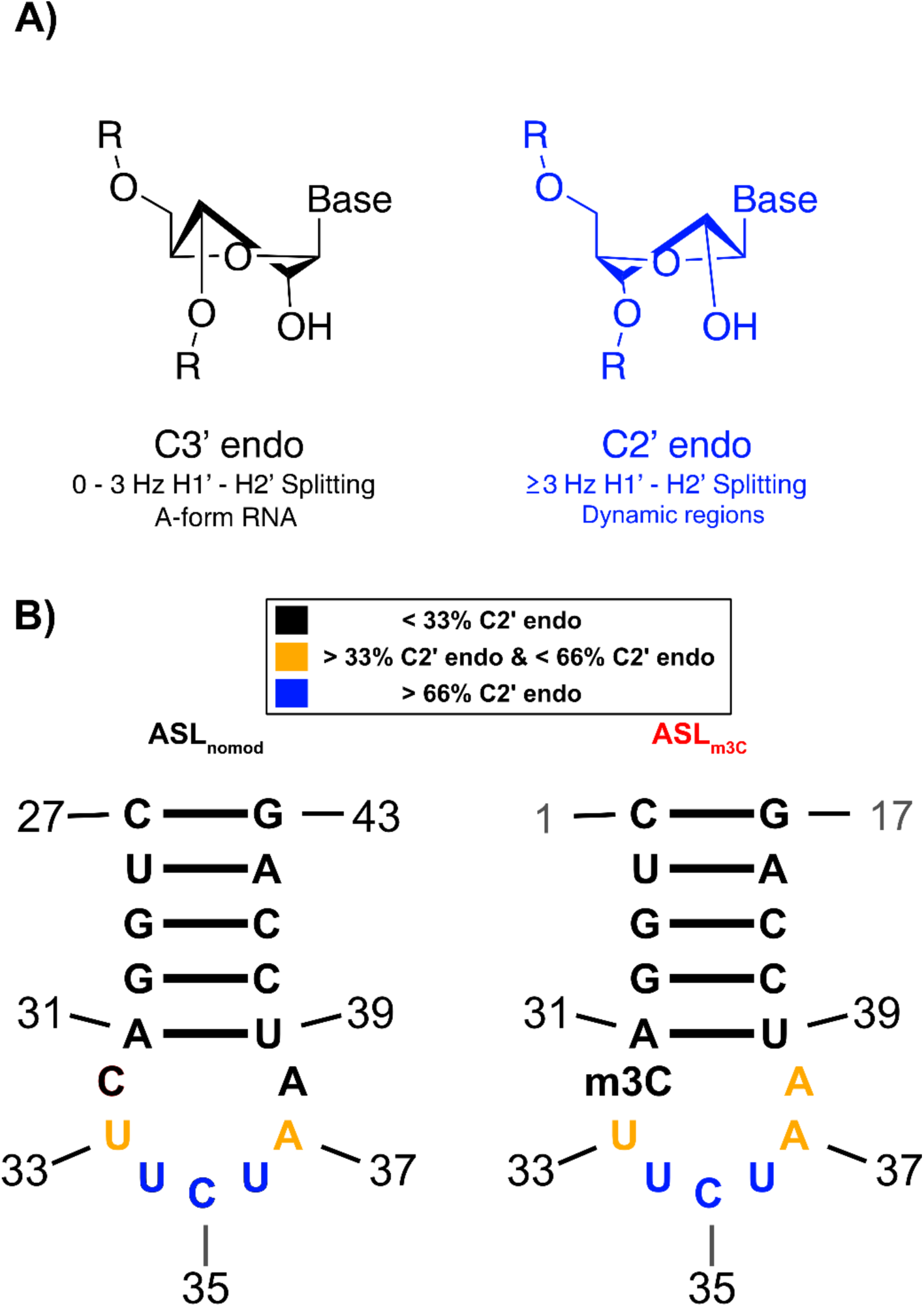
A: The two most prevalent sugar conformations in RNA are C3’ endo (black) and C2’ endo (blue). NMR indicates a preference for C2’ endo when significant splitting is seen between the H1’ and H2’ resonances. B: The sugar conformation preference of the residues within ASLnomod and ASLm3C is shown. Residues preferring C2’ endo are shown in blue whereas residues preferring C3’ endo are shown in black.

**Table 4.** Percentage of C2’ endo conformation.

| **Table 4. Percentage of C2' endo conformation** |  |  |
| --- | --- | --- |
| Residue Number | ASL<sub>nomod</sub> | ASL<sub>m3C</sub> |
| U33 | 53 | 58 |
| U34 | 79 | 71 |
| C35 | 70 | 86 |
| U36 | 65 | 70 |
| A37 | 50 | 36 |
| A38 | 28 | 43 |

### The pairing shape of C32 – A38 is maintained after m3C32 modification

The breadth of structures adopted by C32 – A38 in tRNA structures is wide (Figure S7)[68–70]. Canonical structure for the 32 - 38 pairing within tRNA is regarded as between the H6 aminos of the purine at position 38 and the carbonyl of the pyrimidine at position 32 (Figure S7) [21, 22]. Several lines of evidence in our NMR data suggest a canonical tRNA C32 – A38 pairing in ASL_nomod_ (Figure 5). At short mixing time (75 ms) in ASL_nomod_, the only cross-strand NOE observed in this region is A38 H2 – U33 H1’ (as expected in A-form RNA). At longer mixing time, a relatively weak A38 H2 – C32 H1’ NOE can also be seen, indicating C32 – A38 interaction is likely. There is one hint of hydrogen bonding observed on unmodified C32’s pairing face. C32’s H4 amino protons display two chemical shifts, thus indicating that a hydrogen bond may be present, hindering free rotation of H4 and causing the aminos to be in different environments. As typical A-form NOEs arising from A38 are observed (i.e. to U33 H1’) but no inter-base C32 – A38 NOEs are observed (i.e. amino to adenine H2)[65], one can conclude that the C32-A38 pair in ASL_nomod_ is in an A-form like conformation. Modeling of this pair indicated C32 – A38 is very similar to the canonical tRNA pairing with A38 H6 amino hydrogen bonded to the C32 O2 carbonyl (Figure 5A). In this case, the surrounding NMR derived distances (and the force field) result in stabilization of the canonical pairing during modeling. This modeled structure of the C32 – A38 pairing agrees with the minimal NOE information available regarding C32 – A38, as this pairing puts C32 H4 aminos and A38 H2 far away from one another (no NOE was seen between C32 H4 and A38 H2). Given the weaker nature of the one hydrogen bond C32 – A38 pairing, it is likely that C32 – A38 serves as a transition between the stable stem and dynamic pyrimidine residues in the remainder of the hairpin.

**Figure 5.**
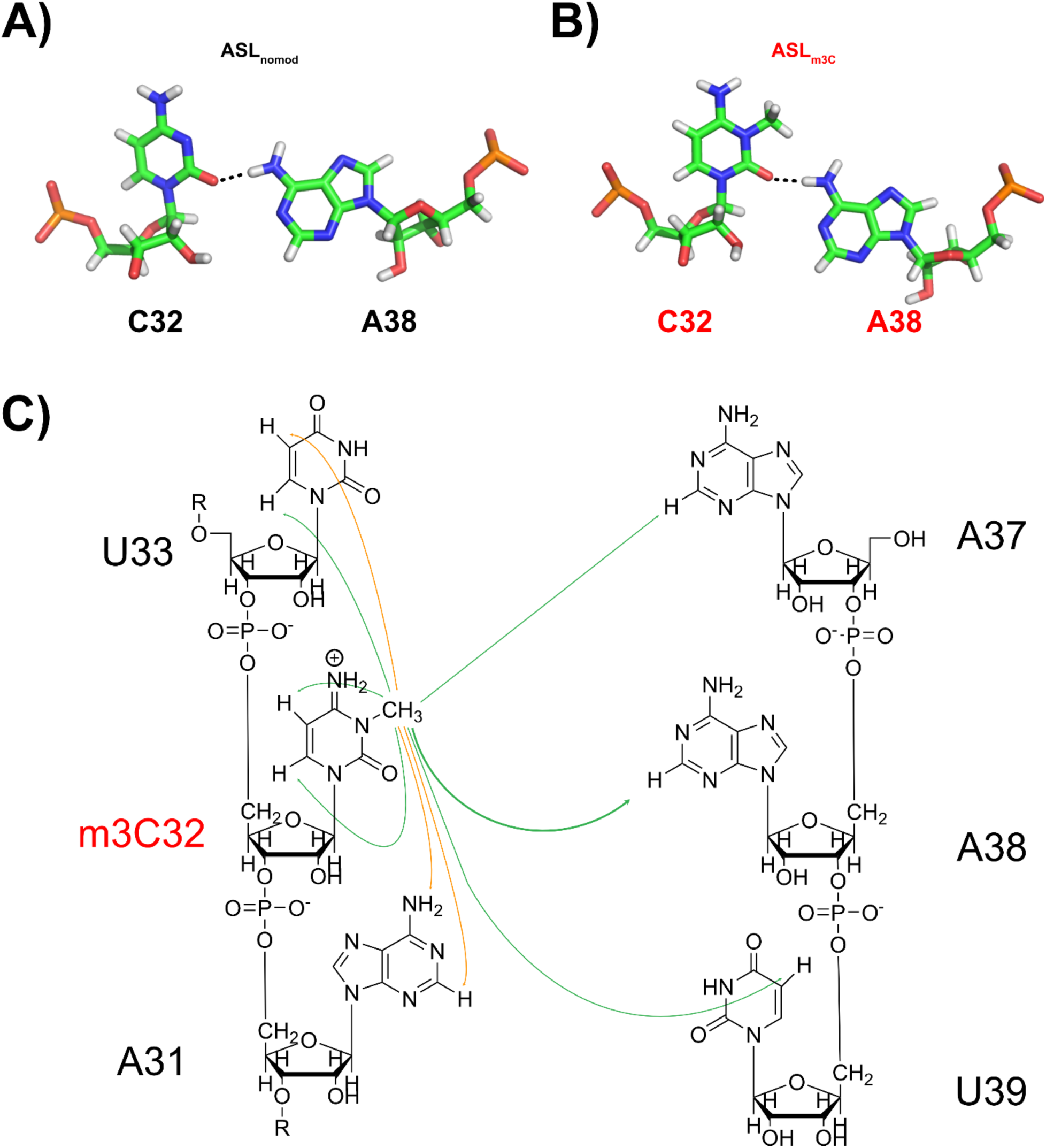
Conformation of C32 – A38 pairing before modification (A) and after m3C modification (B) remains the same despite modification. C: NOEs from the m3C methyl group. NOEs shown in green are only visible at long mixing time (400 ms) whereas NOEs shown in orange are visible at short mixing time.

After m3C32 modification, it is possible to get a better model of C32’s location relative to A38. The m3C32 nucleotide at position 3 of the base has a methyl group on its Watson-Crick pairing face (Figure 1C), thus providing a reporter inside the helix that can inform on m3C32’s position. Nevertheless, only a few strong cross-strand NOEs are seen from m3C32’s methyl group. The only cross-strand NOEs observed from the methyl group are to A37 H2, U39 H5, and A38 H2 (Figure 5C), which are all weak peaks and only observed at longer mixing times. At shorter mixing time, NOEs from the m3C32 methyl are seen to the adjacent U33 H5, A31 H2, and A31 H6 (Figure 5C). The shorter mixing time NOEs indicate a shorter inter-atomic distance, positioning m3C32 squarely within the 31-32-33 base stack. The two previously observed peaks due to C32 aminos (in ASL_nomod_) are not observed with m3C32 because the amino group has been replaced with an imine group. A new resonance appears for the H4 imine with a chemical shift of 9.80 ppm (Figure 1D). NOEs from H4 are to m3C32 itself, suggesting H4 is not participating in pairing. This lack of NOEs across the helix from m3C32’s methyl group (or H4 imine proton) is in line with the lack of direct evidence observed for the C32 – A38 one hydrogen bond pair (Figure 5A). Using the methyl resonances in modeling results in a similar one hydrogen bond pairing between m3C32 and A38 (Figure 5B). This pairing maintains the A38 H6 – m3C32 O2 one hydrogen bond pairing and orients the methyl group so that it is protrudes out into the major groove. The structure of this pairing explains the lack of thermal melting effects for m3C32 modification at higher salt concentrations (Table 1 and 2), as the conformation of m3C32 – A38 is not changing relative to C32 – A38. Furthermore, the pairing provides an explanation for how C32 can be modified while still maintaining the crucial 32 – 38 shape that is likely necessary for successful translation [21, 23, 71, 72]. No evidence of C32 flipping out was observed, such as direct A31 – U33 NOEs. However, this does not necessarily mean that C32 always remains in the helix, because a direct A31 – U33 NOE could not be observed due to overlap.

### m3C modification affects anticodon hairpin loop dynamics and structure

Despite being relatively distant from the modification, changes within the UCU anticodon are observed due to modification (Figure 2 and Figure 6). The primary locations for structural changes due to methylation are U34 and C35. In the absence of methylation, several NOEs indicating dynamic behavior of U34 are observed. U33 H6 - C35 H5, U33 H2’ – C35 H6, and U33 H2’ – C35 H5 are all visible at long mixing times (Figure S8). Evidently, the two non-consecutive C35 / U33 bases can come close. For approach of U33 to C35, U34 is likely extruded some percentage of the time. It is worth noting that all resonances within the anticodon (U34, C35, and U36) have similar chemical shifts prior to modification, as all are dynamic. Thus, some potential NOEs cannot be definitively assigned within the loop due to overlap. Also present prior to modification are NOEs indicating that A37 is near C35 (Figure S6), giving C35 a central position within the hairpin loop while both U34 and U36 have more dynamic positions (Figure 6 and Figure 7A). This is consistent with uracil’s predilection for not stacking [73].

**Figure 6.**
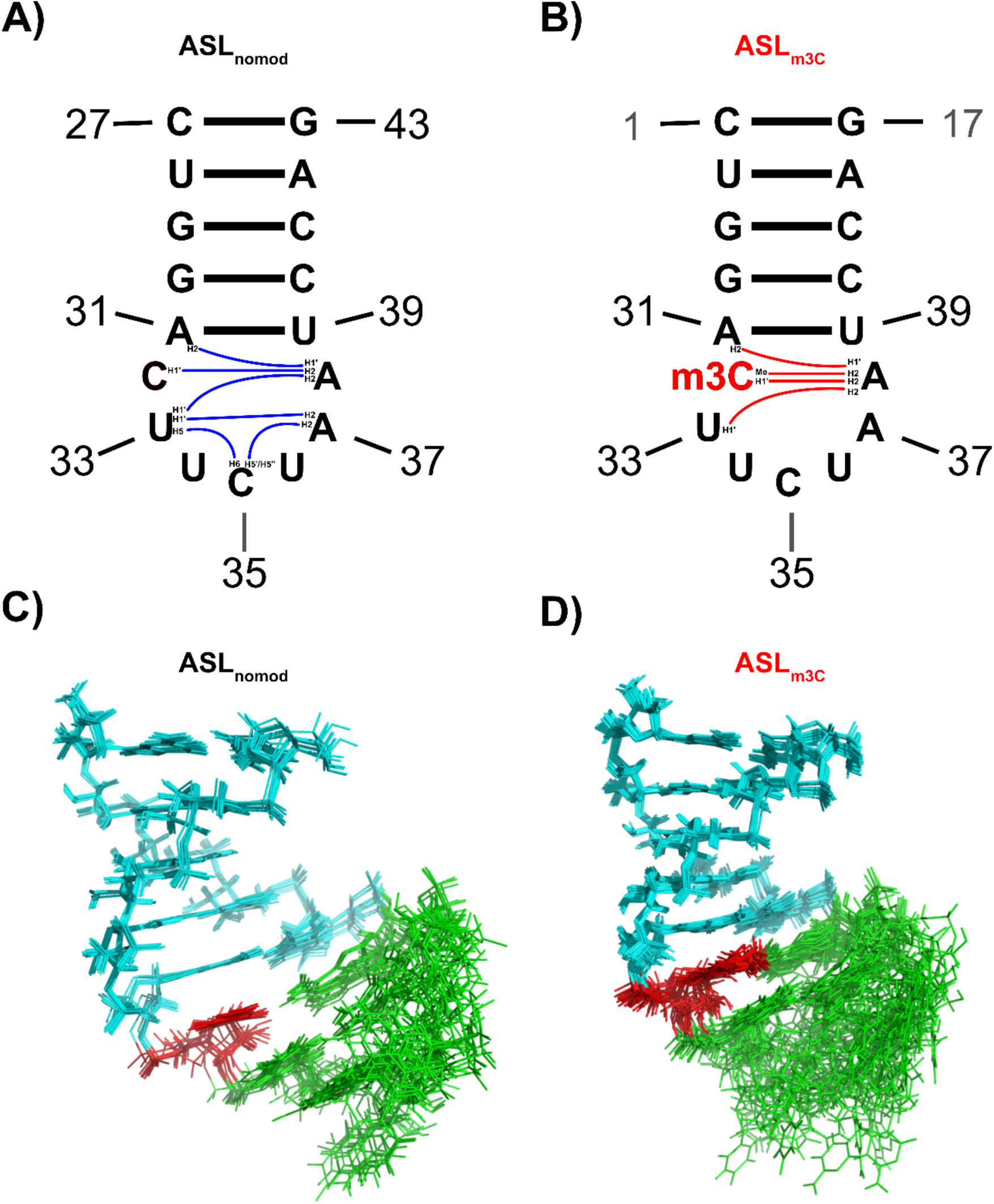
Shown are long range (|I – j| ≥ 2) NOE connections observed within the loops for ASL_nomod_ (A, blue) and ASL_m3C_ (B, red). Shown NOEs include those from both short mixing time (75 ms) and long mixing time (400 ms). Overlaid NMR conformers for ASL_nomod_ (C) and ASL_m3C_ (D) demonstrate increased variability within the loop of ASL_m3C_ due to a lack of long-range NOEs for ASL_m3C_.

**Figure 7.**
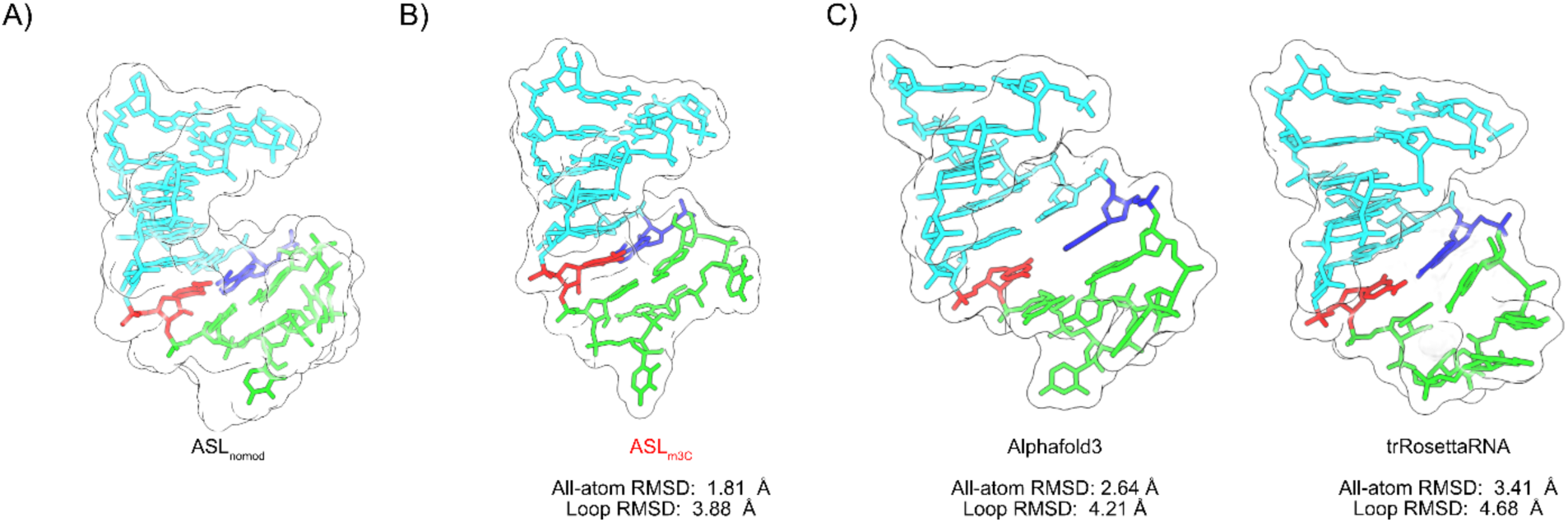
A: Lowest energy NMR conformer of ASL_nomod_. B: Lowest energy NMR conformer of ASL_m3C_ C: Structures of ASL_nomod_ predicted by Alphafold3 and trRosettaRNA. Shown below the predicted structure is the all-atom RMSD and loop RMSD using the lowest energy NMR conformer as a reference. C: Lowest energy ASL_m3C_ structure modeled with NMR-derived restraints. The corresponding RMSD values for ASL_m3C_ used the lowest energy ASL_nomod_ conformer as a reference structure and excluded residue 32 due to the presence of the methyl group.

In the modified ASL_m3C_, all long-range hairpin loop NOEs originating in the anticodon are no longer present, thus indicating a less structured and more dynamic anticodon (Figure 6). In general, dynamics appear to be increased after m3C modification. Evidence for this is provided by a lack of assigned long range NOEs within the anticodon, an increase in C2’ endo character for C35 (Table 4), A38 displaying C2’ endo character, A38 displaying signs of intermediate chemical exchange, and the A31 – U39 pairing destabilization. Altogether, this points towards m3C inducing disorder in the hairpin loop.

Sugar pucker within the anticodon (position 32 through 38) is primarily in the C2’ endo conformation in both unmodified and modified ASLs (Figure 4 and Table 4). H1’-H2’ splittings for residues U33 – U36 are in the 5 – 7 Hz range, indicating prevalence of the C2’ endo sugar pucker. In pure C2’ endo conformation, the distance from H2’ to the H6 of the same residue becomes quite close (r = 2.6 ± 0.5 Å, Table S4). Pure C2’ endo is expected to have a H1’ – H2’ splitting of ~ 7.8 Hz [74]. This starkly contrasts to C3’ endo, where the ribose adopts an H2’ – H6 distance of r =4.0 ± 0.1 Å (Table S4) and H1’ – H2’ splitting of 1.1 Hz for pure C3’ endo [74]. For residues U34, C35, and U36 in both modified and unmodified ASLs, there is a strong n H2’ – n H6 NOEs indicative of C2’ endo behavior. This, combined with strong H1’ – H2’ splitting (Figure 4 and Table 4), and the fact that residues U33 – A37 appear in a short mixing time TOCSY indicate the proportion of bases U33 – U36 adopting C2’ endo sugar pucker is high. Notably, there is an increase in C35’s C2’ endo character after modification, which is in line with C35 becoming more disordered (Table 4). A37 and A38 both display intermediate C2’ endo character in the presence or absence of m3C. For A37, the percentage of C2’ endo (Table 4) is lower when m3C32 is present, perhaps due to due to syn bases preferring a C3’ endo ribose pucker [67]. In ASL_m3C_, residue A38’s sugar adopted an intermediate splitting indicating that it can likely access both C2’ and C3’ endo like behavior. Additionally, A38 in ASL_m3C_ displays some temperature sensitivity not seen in the remainder of the residues, showing signs of intermediate exchange at lower temperatures. This intermediate behavior of A38 is further suggested by a sharpening of the A38 resonances at higher temperatures. In unmodified C32 or m3C32, there is no indication of C2’ endo behavior.

### Overall structural details for ASL_nomod_ and ASL_m3C_

Between ASL_nomod_ and ASL_m3C_, NMR conformers show differences in disorder within the hairpin loop (Figure 6). ASL_nomod_ structure is well determined within the hairpin loop due to the previously discussed C35 NOEs to A37 and U33 (Figure 6). However, the anticodon region within ASL_m3C_ is underdetermined due to a lack of inter-residue NOEs. Thus, the aligned conformers of ASL_m3C_ show considerable structural dispersion within the anticodon region and a higher RMSD of the conformers within the loop (Figure 6 and Table 5). This lack of NOE information within ASL_m3C_ anticodon region is likely reflective of significant dynamics within the region.

**Table 5.** Structural modeling constraints and statistics.

| **Table 5. Structural modeling constraints and statistics** |  |  |
| --- | --- | --- |
| Number of restraints | ASL<sub>nomod</sub> | ASL<sub>m3C</sub> |
| All distance restraints | 277 | 232 |
| All NOE restraints | 255 | 210 |
| Intra-nucleotide | 175 | 134 |
| Inter-nucleotide | 102 | 76 |
| Long range | 50 | 44 |
| Hydrogen bond | 21 | 21 |
| Backbone dihedral | 76 | 76 |
| RMSD of conformers (Å) |  |  |
| All | 1.03 | 3.13 |
| Stem | 0.40 | 0.45 |
| Loop | 1.52 | 4.86 |
| Noe violations |  |  |
| Distance (> 0.1 Å) | 0 | 0 |
| Dihedral Angle (>5 °) | 0 | 0 |

The U-turn motif within the ASL has been theorized to be important for binding of tRNA by the ribosome [75]. The motif has a clear NMR signature: an NOE is seen between U33 H1’ and the n+2 residue’s H6/H8, an additional imino peak is observed for the U33 imino proton due to stable interaction with the 35-P-36 phosphate, a shift in phosphate chemical shift for the U33-P-U34 phosphate, and an adoption of C3’ endo sugar conformation within the 34-35-36 stack [17]. However, none of these NMR-based signatures are observed in either the unmodified or m3C32 modified ASL, indicating that the U-turn like motif is not adopted in either case. The U-turn motif results in a stack of the next 3 residues (34, 35 and 36) after the turning U33. Given that uracils are generally considered to stack poorly [73] and that the similar tRNA-Lys-UUU requires U34 modification to adopt a U-turn like structure, it is unsurprising to see the structure not adopt a U-turn. Apparently, m3C modification is not sufficient to induce formation of the canonical U-turn structure. However, it should be noted that a possible U33 H1’ – C35 H6 NOE cannot be seen due to overlap of U33 H1’ and C35 H5.

### Comparison of ASL_nomod_ to 3D predicted structure

Several predictions of unmodified ASL structure were compared with the unmodified structure determined from experimental data (Figure 7). Predictions from AlphaFold3 [52] and trRosettaRNA [53] were compared to the unmodified ASL structure by all-atom RMSD and loop specific (residues 32-38) RMSD. In addition, several structural features of ASL_nomod_ that were clearly indicated by NMR data were compared with features within the predicted structures (Table 6). Notably, several features observed in our experimental data were not accurately reflected by the predicted structures (Figure 7). In general, Alphafold3 performed the best, reflecting many structural features consistent with the NMR data and providing a model that closely resembles the ASL_nomod_ structure. This aligns with recent reports indicating that AlphaFold3 can exhibit surprising accuracy in predicting RNA-only structures [76]. Due to the existence of a modification within ASL_m3C_, no predictions were made for ASL_m3C_, but ASL_m3_C is shown within Table 6 and Figure 7 so that its structural features may be easily compared to those of ASL_nomod_.

**Table 6.** Comparison of predicted structures and ASL_m3C_ to ASL_nomod_.

| **Table 6. Comparison of predicted structures and ASL<sub>m3C</sub> to ASL<sub>nomod</sub>** |  |  |  |  |
| --- | --- | --- | --- | --- |
| Feature<sup>a</sup> | ASL<sub>nomod</sub> | AlphaFold3 | tr Rosetta RNA | ASL<sub>m3C</sub> |
| Sugar pucker<sup>b</sup> C32 | C3' endo | **C3' endo** | **C3' endo** | **C3' endo** |
| Sugar pucker<sup>b</sup> U33 | C2' endo<sup>c</sup> | C3' endo<sup>c</sup> | **C2' endo<sup>c</sup>** | **C2' endo<sup>c</sup>** |
| Sugar pucker<sup>b</sup> U34 | C2' endo | **C2' endo** | Non C2'/C3' endo | **C2' endo** |
| Sugar pucker<sup>b</sup> C35 | C2' endo | **C2' endo** | Non C2'/C3' endo | **C2' endo** |
| Sugar pucker<sup>b</sup> U36 | C2' endo | **C2' endo** | Non C2'/C3' endo | **C2' endo** |
| Sugar pucker<sup>b</sup> A37 | C2' endo<sup>c</sup> | C3' endo<sup>c</sup> | Non C2'/C3' endo<sup>c</sup> | C3' endo<sup>c</sup> |
| Sugar pucker<sup>b</sup> A38 | C3' endo<sup>c</sup> | **C3' endo<sup>c</sup>** | **C3' endo<sup>c</sup>** | **C3' endo<sup>c</sup>** |
| C32 - A38 pairing | 1 H-bond<br>C32 O2 – A38 H6 | No pair | No pair | **1 H-bond**<br>**C32 O2 – A38 H6** |
| C35 | Central: U33,<br>U34, U36, A37<br>surround C35 | **Central: U33,**<br>**U34, U36, A37**<br>**surround C35** | Flipped | Flipped |
| RMSD (all-atom) | - | 2.64 Å | 3.41 Å | 1.81 Å<sup>d</sup> |
| RMSD (loop) | - | 4.21 Å | 4.68 Å | 3.88 Å<sup>d</sup> |
| <sup>a</sup>Agreement with ASL<sub>nomod</sub>'s NMR data is indicated by bold<br><sup>b</sup>Sugar pucker deemed to be C2' endo if ≥ 50% is C2' endo. If < 50% C2' endo, deemed to be C3' endo<br><sup>c</sup>Indicates a residue with intermediate C2'/C3' endo character indicated by NMR (less than 65% endo character).<br><sup>d</sup>RMSD calculated to all residues besides modified residue |  |  |  |  |

## Discussion

Here, we show that m3C modification of the ASL of tRNA-Arg-UCU-4-1 shifts the structure of the hairpin loop and adjacent A31 – U39 pairing, while maintaining the C32-A38 pairing’s canonical shape. This change in structure suggests that upon m3C32 modification, the tRNA ASL changes in ways that may impact function. Overall, our study reveals how the m3C modification can alter the dynamics and structure of the uniquely expressed tRNA-Arg-UCU-4-1 isodecoder and other tRNAs containing m3C in the ASL.

In the context of tRNA-Arg-UCU-4-1, m3C32 modification shifts the structure of the hairpin loop and nearby stem (Figure 2). This shift in structure results in adoption of elevated syn base character in A37 (Figure 3), increased dynamics within the hairpin loop (Figures 5 and 6), and a decrease in A31-U39 pairing stability (Figure 1D). These changes could affect tRNA interaction with other tRNA modification enzymes, aminoacyl tRNA synthetases, and the ribosome. The structures of tRNA-Arg-UCU-4-1’s ASL in both unmodified and m3C modified states (Figure 6) likely represent freeze frames of intermediates along the path to a fully modified tRNA [77]. It is likely that the changes in structure described herein shift how tRNA-Arg ASLs can interact with its various binding partners, thus guiding the tRNA towards becoming a functional tRNA.

Interestingly, m3C32 modification appears to only subtly affect the C32 – A38 pairing. Structural modeling (Figure 5) of the modified m3C32 – A38 base pair indicated that the orientation of the pairing is only subtly shifted. After modification, a similar A38 H6 – C32 O2 interaction is observed. The m3C32 methyl group thus orients itself facing out of the major groove (Figure 5B), potentially allowing for altered interactions through the major groove. Similarity of the unmodified pairing to the m3C32 modified pairing is likely required by evolution, as the shape of the 32 – 38 pairing is reportedly important for maintaining tRNA’s role in translation [23, 71, 72, 78]. Thus, maintenance of the isosteric nature of the 32 – 38 pair is likely to be important, as mere reversal of the pairing may hinder ribosomal interaction [21, 72].

Modification of m3C32 within tRNA-Arg-UCU-4-1 has minor impact upon the thermodynamics of the hairpin loop (Table 1). This is interesting, given that replacement of C with m3C in duplexes or single nucleotide mismatches has been shown to be strongly destabilizing in all instances tested (including in C-A mismatches)[24]. In the specific instance of replacing a C with m3C at position 32 in tRNA, modification may represent a way of ensuring the shape of the C32 – A38 pairing (Figure 5). m3C modification at position 32 will not only promote correct C32 – A38 pairing orientation but will also discourage alternative base pairing schemes in which C32 interacts with a residue other than A38. Thus, we hypothesize that m3C32 modification helps to ensure faithful translation by discouraging base pair structures other than the canonical 32 – 38 structure.

A canonical U-turn is beneficial for tRNA binding to the ribosome [16, 19, 79, 80]. There was no NMR based evidence of U-turn formation in either modified or unmodified tRNA-Arg-UCU-4-1 ASL structures. Furthermore, the anticodon residues showed significant C2’ endo ribose pucker (Figure 4) rather than C3’ endo pucker that would be expected if the residues were stacked in a U-turn [19]. Increased C2’ endo conformation has been shown to reduce affinity for the ribosome [16, 19, 20, 80]. Thus, it is likely that tRNA-Arg-UCU-4-1 may have reduced ribosomal binding in the absence of further modification. In addition, it has been shown previously that U34 modifications increase the residue 34-35-36 stack within the similar tRNA-Lys-UUU [19] and increase ribosomal affinity [16]. t6A modification at position 37 in tRNA-Lys-UUU has also been shown to modulate ribosome binding in tRNA-Lys-UUU [16]. Therefore, m3C modification may represent one of the first modifications made upon the ASL to help ensure the correct ASL configuration is attained. Further modifications, such as t6A37 or mcm5s2U34, can then push the structures towards higher affinity ribosomal interaction.

Here we show that m3C32 modification stabilizes the effects of negatively charged backbone phosphates by introduction of a positive charge on m3C32’s base (Table 2). Previous thermodynamic studies of m3C modifications have demonstrated that substituting m3C for C in any C – N pairing (where N is A, C, G or U) is strongly destabilizing [24]. Strong destabilization, such as observed by Mao et al., implies a corresponding change in structure. C32 pre-exists in a loop region and modification results in minimal change thermodynamically (Table 1). This lack of change is interesting, given that m3C modification appears to destabilize A31 – U39 (Figure 1D) and induce dynamics within the hairpin loop (Figure 4). In the case of tRNA modification at position 32, the enthalpic destabilization induced by m3C modification is likely offset by its electrostatic stabilization and entropic effects.

Previous studies have reported the existence of a pH dependent protonated C_32_ − A^+^_38_ pair with a pKa of approximately 6 for tRNA-Lys-UUU [58]. tRNA-Arg-UCU-4-1 has a very similar sequence within the ASL, thus its stabilization at low pH is also likely due to C_32_ − A^+^_38_ (Table 3). Since NMR was carried out with a pH of 6.3 (above the approximate C_32_ − A^+^_38_ pKa), it is unsurprising to see little evidence of C_32_ − A^+^_38_. Despite the acidic pKa of C_32_ − A^+^_38_, it is still possible that a protonated C32 – A^+^_38_ pairing is biologically relevant in the unmodified tRNA-Arg-UCU and tRNA-Lys-UUU. Acidic pH changes approaching pH 6 are common detected during inflammation [81], ischemic stroke [82], myocardial ischemia [83], and many cancers [84]. Such a change in tRNA structure could correspond to a subsequent shift in tRNA function/translation at lower pH’s for tRNAs that contain a C32 – A38 pairing.

It is possible that m3C modification impacts other modification circuits elsewhere in the tRNA body by modulating tRNA ASL structure. In this light, the A37 structural shift (Figure 3) becomes especially relevant given that A37 in tRNA-Arg-UCU-4-1 can be modified with t6A. This t6A modification, within the very similar tRNA-Lys-UUU ASL, was reported to cause structural rearrangement [17]. If both modifications individually cause reorganization, it is likely that the ASL’s structure will adopt a different orientation if both modifications are present. Crosstalk between the m3C and t6A modifications has already been noted in the case of tRNA-Thr, where t6A modification alone allows tRNA to be modified by METTL2A [85]. Furthermore, prior t6A modification can help promote m3C modification by METTL8 in mitochondria [25]. Given that the t6A and m3C have opposite formal charges upon their bases, crosstalk in the opposite direction (i.e. m3C promoting t6A modification) seems probable.

## Abbreviations

ASL: Anticodon stem-loop
m3C: 3-methylcytosine
t6A: N6-threonylcarbamoyladenosine
mcm5s2U: methoxycarbonylmethyl thiouridine
NOESY: Nuclear Overhauser spectroscopy
NOE: nuclear Overhauser effect
TOCSY: total correlation spectroscopy
HSQC: hetero single quantum coherence
RESP: restrained electrostatic potential;

## Acknowledgements

NMR data was collected at SUNY ESF. We acknowledge the upgrade of the 600 MHz NMR spectrometer at SUNY-ESF under NSF grant CHE-1048516; and the acquisition of the 800 MHz NMR spectrometer through NIH grant S10 OD012254. This study made use of NMRbox: National Center for Biomolecular NMR Data Processing and Analysis, a Biomedical Technology Research Resource (BTRR), which is supported by NIH grant P41GM111135 (NIGMS) [86]. Structural analysis used ChimeraX. ChimeraX was developed by the Resource for Biocomputing, Visualization, and Informatics at the University of California, San Francisco, with support from National Institutes of Health R01-GM129325 and the Office of Cyber Infrastructure and Computational Biology, National Institute of Allergy and Infectious Diseases. Graphical abstract contains a tRNA image created in BioRender (https://BioRender.com/t33r041).

We thank Doug Turner, Scott Kennedy, and the members of the Fu lab for reading and providing useful feedback regarding the manuscript. Additionally, we thank Charlie Fry at SUNY ESF’s Analytical and Technical Services (ACTS) who aided in NMR data acquisition. We also thank the lab of Benjamin Miller at the University of Rochester Medical Center for generous access to their Shimadzu UV-1800 spectrophotometer.

## Accession numbers

NMR conformers for unmodified and m3C modified ASLs are deposited in the PDB with accession codes 9ECQ and 9ECR, respectively. NMR shifts and assignments for the unmodified and m3C modified ASLs are available at the BMRB with codes 31216 and 31217, respectively.

## Author contributions: CRediT

**Kyle D Berger:** Conceptualization, Data curation, Formal Analysis, Investigation, Methodology, Software, Visualization, Writing – original draft, Writing – review and editing. **Anees Mohammed**: Formal Analysis, Methodology, Software, Writing – review and editing. **David H Mathews**: Funding acquisition, Project Administration, Resources, Supervision, Writing – review and editing. **Dragony Fu**: Conceptualization, Funding acquisition, Project Administration, Resources, Supervision, Writing – original draft, Writing – review and editing.

## Funding sources

This work was funded by NIH R01 GM143145 to D.F. and NIH R35GM145283 to D.H.M.

## Notes

### Competing Interest Statement

The authors have declared no competing interest.

