## Supplemental Information for "Structural impact of 3-methylcytosine modification on the anticodon stem of a neuronally-enriched arginine tRNA"

$$\Delta G_{37}^{^{\circ}}\left( {ASL}_{nomod} \right)= G_{37}^{^{\circ}}\left( Stem \right)+G_{37}^{^{\circ}}\left( Terminal mismatch \right)+G_{37}^{^{\circ}}\left( Hairpin \right)$$

$$\Delta G_{37}^{^{\circ}}\left( {ASL}_{nomod} \right)= G_{37}^{^{\circ}}\left( CG followed by UA \right)+G_{37}^{^{\circ}}\left( UA followed by GC \right)+G_{37}^{^{\circ}}\left( GC followed by GC \right)+G_{37}^{^{\circ}}\left( GC followed by AU \right)+G_{37}^{^{\circ}}\left( AU end penalty \right)+G_{37}^{^{\circ}}\left( AU followed by CA \right)+G_{37}^{^{\circ}}\left( 7 nt hairpin \right)$$

$$\Delta G_{37}^{^{\circ}}\left( {ASL}_{nomod} \right)=\left( -2.1 kcal/mol \right)+\left( -2.1 kcal/mol \right)+\left( -3.3 kcal/mol \right)+\left( -2.4 kcal/mol \right)+\left( 0.45 kcal/mol \right)+\left( -0.6 kcal/mol \right)+\left( 6.0 kcal/mol \right)$$

$$\Delta G_{37}^{^{\circ}}\left( {ASL}_{nomod} \right)=-4.05 kcal/mol$$

Figure S1. Calculation of folding free energy for tRNA-Arg-UCU-4-1 ASL_nomod_. Calculation used 2004 RNA ΔG°_37_ parameters.

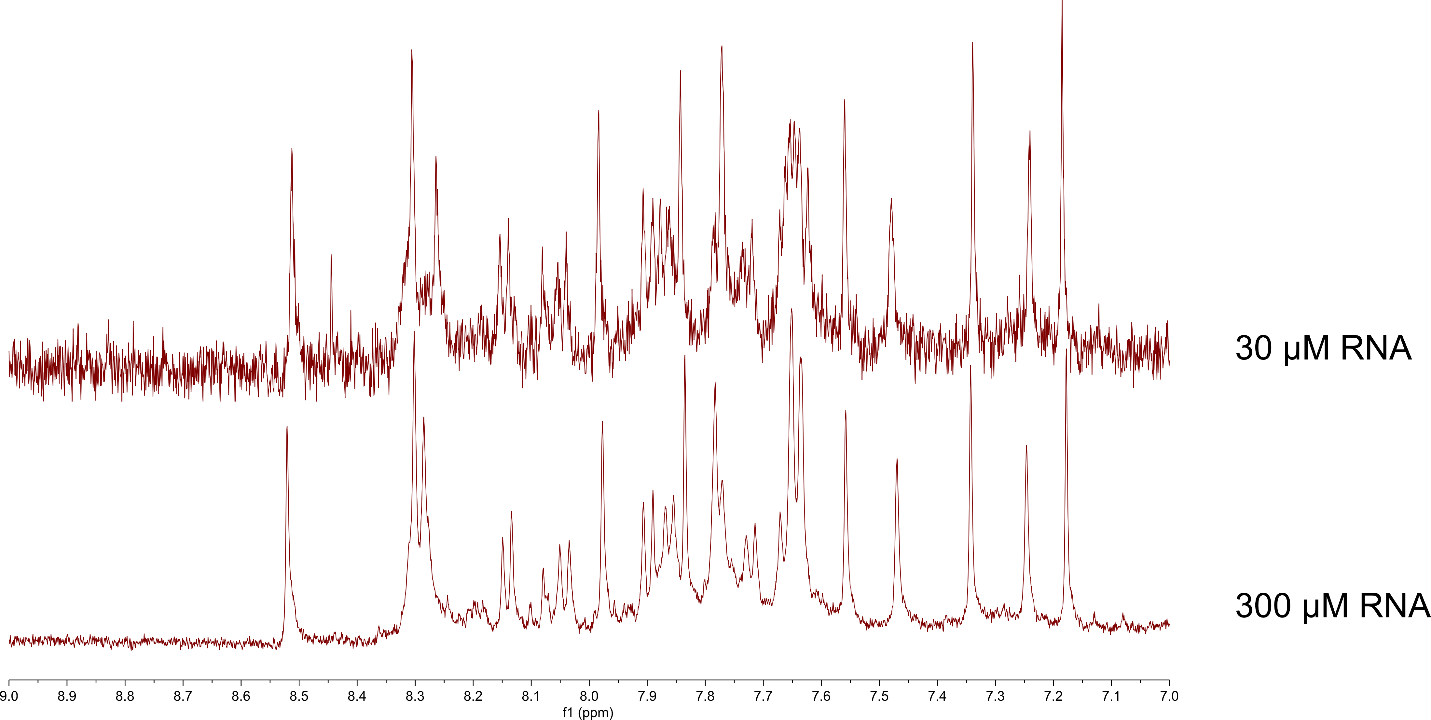

Figure S2. 1D ASL_nomod_ spectra at two different RNA concentrations

Representative 1M NaCl Melt curves

Absorbance (normalized)

| Temp (C) | m3C |
| --- | --- |
| 19.8 | 0 |
| 21.2 | 0.00990099 |
| 22.3 | 0.019802 |
| 23.3 | 0.029703 |
| 24.2 | 0.029703 |
| 25.4 | 0.019802 |
| 26.4 | 0.029703 |
| 27.6 | 0.029703 |
| 28.8 | 0.039604 |
| 30.3 | 0.039604 |
| 31.1 | 0.039604 |
| 31.9 | 0.049505 |
| 32.7 | 0.049505 |
| 33.9 | 0.049505 |
| 34.7 | 0.049505 |
| 35.5 | 0.049505 |
| 36.1 | 0.049505 |
| 37.2 | 0.039604 |
| 38.5 | 0.039604 |
| 39.2 | 0.039604 |
| 40.4 | 0.049505 |
| 41.2 | 0.049505 |
| 42.3 | 0.049505 |
| 43.3 | 0.049505 |
| 44.3 | 0.0693069 |
| 45.3 | 0.0693069 |
| 46.3 | 0.0693069 |
| 47.4 | 0.0792079 |
| 48.3 | 0.0792079 |
| 49.3 | 0.0891089 |
| 50.2 | 0.0891089 |
| 51.3 | 0.108911 |
| 52.5 | 0.118812 |
| 53.7 | 0.128713 |
| 54.8 | 0.138614 |
| 55.7 | 0.148515 |
| 56.8 | 0.168317 |
| 57.6 | 0.178218 |
| 58.6 | 0.19802 |
| 59.5 | 0.217822 |
| 60.4 | 0.227723 |
| 61.2 | 0.257426 |
| 62.3 | 0.277228 |
| 63.3 | 0.306931 |
| 64.4 | 0.346535 |
| 65.2 | 0.366337 |
| 66.3 | 0.39604 |
| 67.2 | 0.445545 |
| 68.3 | 0.475248 |
| 69.3 | 0.514851 |
| 70.4 | 0.564356 |
| 71.5 | 0.594059 |
| 72.3 | 0.623762 |
| 73.2 | 0.663366 |
| 74.2 | 0.693069 |
| 75.3 | 0.732673 |
| 76.3 | 0.762376 |
| 77.3 | 0.782178 |
| 78.2 | 0.80198 |
| 79.3 | 0.831683 |
| 80.3 | 0.841584 |
| 81.3 | 0.861386 |
| 82.3 | 0.881188 |
| 83.1 | 0.891089 |
| 84.3 | 0.90099 |
| 85.4 | 0.910891 |
| 86.2 | 0.920792 |
| 87.2 | 0.930693 |
| 88.3 | 0.940594 |
| 89.5 | 0.940594 |
| 90.6 | 0.960396 |
| 91.8 | 0.960396 |
| 92.6 | 0.970297 |
| 93.9 | 0.970297 |
| 94.8 | 0.970297 |
| 96 | 0.980198 |
| 97.2 | 0.990099 |
| 98.2 | 0.990099 |
| 99.2 | 1 |

| Temp (C) | nomod |
| --- | --- |
| 19.9 | 0 |
| 21.2 | 0 |
| 22.3 | 0 |
| 23.2 | 0.00806452 |
| 24.2 | 0.016129 |
| 25.2 | 0.0241935 |
| 26.2 | 0.0241935 |
| 27.2 | 0.0241935 |
| 28.2 | 0.0322581 |
| 29.2 | 0.0403226 |
| 30.2 | 0.0483871 |
| 31.2 | 0.0483871 |
| 32.2 | 0.0564516 |
| 33.2 | 0.0645161 |
| 34.2 | 0.0725806 |
| 35.2 | 0.0725806 |
| 36.2 | 0.0806452 |
| 37.2 | 0.0806452 |
| 38.2 | 0.0887097 |
| 39.2 | 0.0967742 |
| 40.4 | 0.0967742 |
| 41.1 | 0.0967742 |
| 42.2 | 0.0967742 |
| 43.3 | 0.112903 |
| 44.3 | 0.112903 |
| 45.3 | 0.129032 |
| 46.3 | 0.137097 |
| 47.3 | 0.137097 |
| 48.3 | 0.153226 |
| 49.2 | 0.16129 |
| 50.2 | 0.169355 |
| 51.2 | 0.177419 |
| 52.3 | 0.177419 |
| 53.3 | 0.201613 |
| 54.2 | 0.201613 |
| 55.2 | 0.209677 |
| 56.2 | 0.225806 |
| 57.2 | 0.233871 |
| 58.3 | 0.258065 |
| 59.2 | 0.274194 |
| 60.2 | 0.298387 |
| 61.3 | 0.322581 |
| 62.2 | 0.346774 |
| 63.3 | 0.387097 |
| 64.3 | 0.419355 |
| 65.2 | 0.451613 |
| 66.2 | 0.491935 |
| 67.2 | 0.532258 |
| 68.3 | 0.572581 |
| 69.3 | 0.620968 |
| 70.2 | 0.653226 |
| 71.3 | 0.701613 |
| 72.3 | 0.741935 |
| 73.3 | 0.782258 |
| 74.2 | 0.814516 |
| 75.3 | 0.83871 |
| 76.2 | 0.862903 |
| 77.2 | 0.895161 |
| 78.3 | 0.903226 |
| 79.2 | 0.927419 |
| 80.2 | 0.943548 |
| 81.1 | 0.951613 |
| 82.2 | 0.959677 |
| 83.2 | 0.975806 |
| 84.3 | 0.967742 |
| 85.2 | 0.975806 |
| 86.2 | 0.983871 |
| 87.2 | 0.983871 |
| 88.4 | 0.991935 |
| 89.3 | 0.991935 |
| 90.2 | 0.991935 |
| 91.2 | 0.991935 |
| 92.3 | 1 |
| 93.3 | 1 |
| 94.2 | 1 |
| 95.2 | 1 |
| 96.3 | 1 |
| 97.3 | 0.983871 |
| 98.2 | 1 |
| 99.3 | 0.991935 |
| 100 | 0.991935 |

Figure S3. Representative overlaid normalized melting data for ASL_nomod_ and ASL_m3C_

$$\Delta G_{37}^{^{\circ}}\left( ASL \right)= G_{37}^{^{\circ}}\left( Stem \right)+G_{37}^{^{\circ}}\left( Terminal mismatch \right)+G_{37}^{^{\circ}}\left( Hairpin \right)$$

$$AG_{37}^{^{\circ}}\left( Terminal mismatch \right)+G_{37}^{^{\circ}}\left( Hairpin \right)=\Delta G_{37}^{^{\circ}}\left( {ASL}_{meas} \right)-G_{37}^{^{\circ}}\left( {Stem}_{pred} \right)$$

$$AG_{37}^{^{\circ}}(HP+TM)=\Delta G_{37}^{^{\circ}}\left( {ASL}_{meas} \right)-G_{37}^{^{\circ}}\left( {Stem}_{pred} \right)$$

Figure S4. Calculation of $G_{37}^{^{\circ}}(HP+TM)$ to estimate hairpin loop thermodynamic contributions to folding

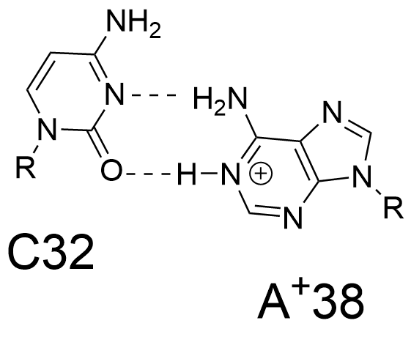

Figure S5. Structure of C32 – A^+^38 pairing

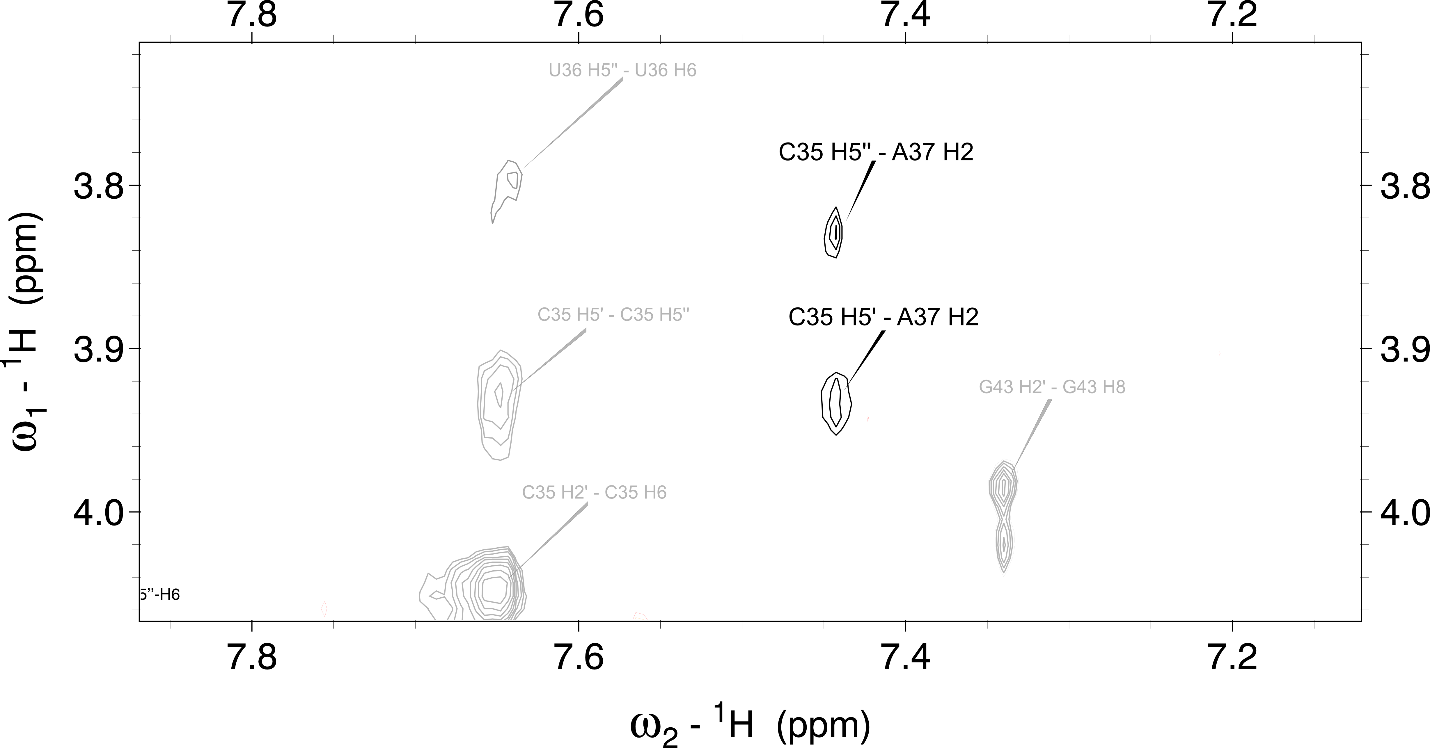

Figure S6. Long range inter-nucleotide NOE at short mixing time (75 ms) between C35 and A37.

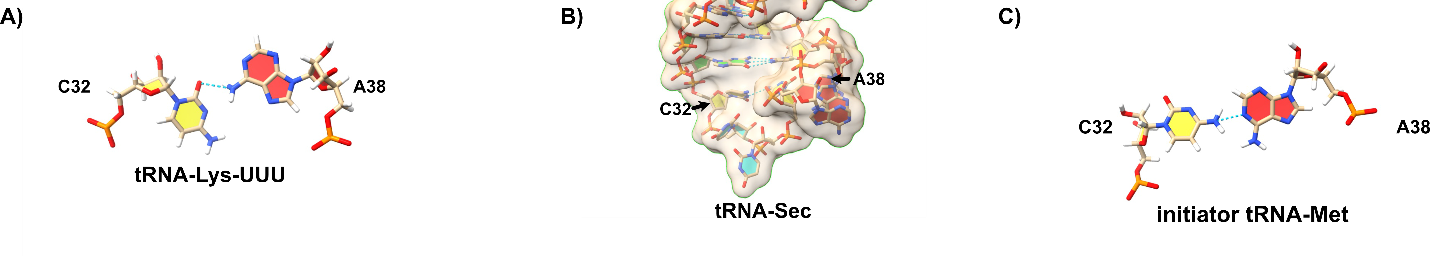

Figure S7. C32 – A38 pairing structures within tRNA

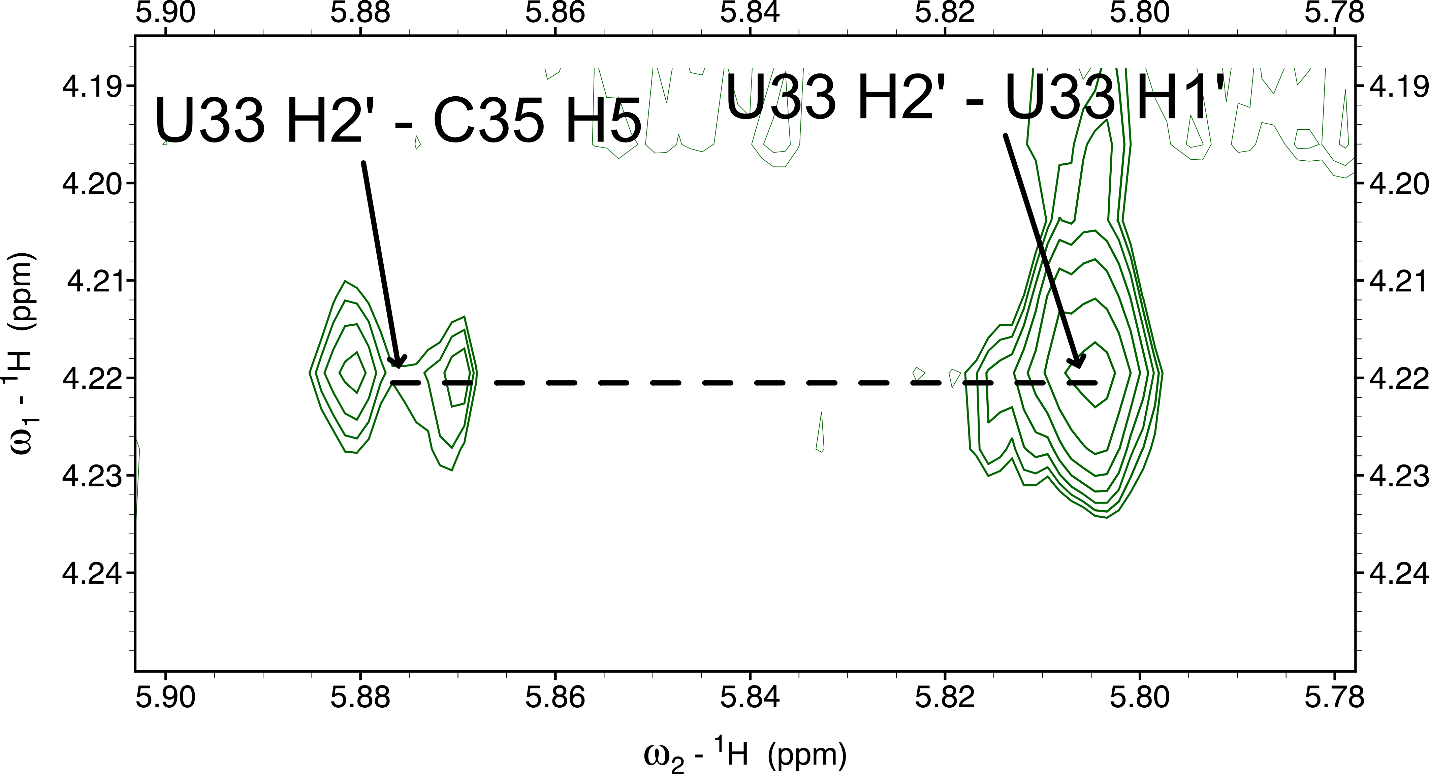

Figure S8. Long range inter-nucleotide NOE at short mixing time (75 ms) between U33 and C35

| Table S1. Melting temperature dependence upon RNA concentration | | |
| --- | --- | --- |
|  | **[RNA] (μM)** | **T_M_** |
| ASL_nomod_ | 1.00 | 70.5 |
|  | 2.00 | 68.0 |
|  | 3.19 | 70.2 |
|  | 5.64 | 69.8 |
|  | 7.41 | 69.9 |
| ASL_m3C_ | 1.77 | 70.5 |
|  | 3.53 | 68.6 |
|  | 4.36 | 68.6 |
|  | 5.37 | 69.4 |

| Table S2. Calculation of Average n H1’ – (n+1) H8 distances for adenines | | | |
| --- | --- | --- | --- |
| **PDB Code** | **Ade 1 (residue #)** | **Ade 2 (residue #)** | **r (n A H1' - n+1 A H8) (Å)** |
| 157D | 5 | 6 | 5.1 |
| 1EVV | 66 | 67 | 4.8 |
| 1K9W | 21 | 22 | 4.8 |
| 1QC0 | 131 | 132 | 5.1 |
| 1SDR | 2 | 3 | 5 |
| 1X8W | 230 | 231 | 5.2 |
| 4XNR | 16 | 17 | 4.4 |
| 4XNR | 29 | 30 | 5.2 |
| 4Y1I | 3 | 4 | 4.7 |
| 4Y1I | 29 | 30 | 4.7 |
|  |  | **AVG:** | 4.9 |
|  |  | **STDEV:** | 0.262466929 |
|  |  | **N:** | 10 |
|  |  | **SE:** | 0.082999331 |

| Table S3. Calculation of Average n H1’ – (n+1) H8 distances for adenines | | | | | |
| --- | --- | --- | --- | --- | --- |
| **Anti** | | | **Syn** | | |
| **PDB Xtal Structure Code** | **Ade #** | **Dist (Å) (H1' - H8)** | **PDB Xtal Structure Code** | **Ade #** | **Dist (Å) (H1' - H8)** |
| 1I9V | A5 | 3.7 | 1F1T | 30 | 2.5 |
| 1DDY | A420 | 3.8 | 1GRZ | 187 | 2.4 |
| 1O3Z | A21 | 3.8 | 5DH8 | 20 | 2.5 |
| 1SDR | A16 | 3.8 | 4ZNP | 29 | 2.7 |
| 2EES | A52 | 3.9 | 4ZNP | 55 | 2.4 |
| 2GDI | A35 | 3.7 | 5C7U | 23 | 2.5 |
| 2OEU | A34 | 3.7 | 5BTP | 48 | 2.6 |
| 3DIG | A47 | 3.7 | 4XNR | 23 | 2.5 |
| 3GX2 | A70 | 3.8 | 4XNR | 64 | 2.4 |
| 4R4V | A746 | 3.6 | 4WCP | 41 | 2.4 |
| 4XNR | A55 | 3.7 | 4R4V | 621 | 2.6 |
| 4Y1I | A75 | 3.8 | 47B0 | 9 | 2.4 |
| 4YB0 | A14 | 3.8 | 4Y1I | 42 | 2.7 |
| 4ZNP | A33 | 3.8 | 4WFL | 44 | 2.5 |
| 5BTP | A18 | 3.7 | 4TZY |  | 2.5 |
|  | **AVG** | 3.75 |  | **AVG** | 2.51 |
|  | **STDEV** | 0.07 |  | **STDEV** | 0.10 |
|  | **N:** | 15 |  | **N:** | 15 |

| Table S4 Calculation of average n H2’ – n H6 distances as a function of whether the base adopts C2’ endo or C3’ endo | | | | | | | |
| --- | --- | --- | --- | --- | --- | --- | --- |
| **Uracil** | | | | | | | |
| **C2’ Endo** | | | | **C3’ Endo** | | | |
| **PDB structure** | **Nucleotide** | **H2'-H6 distance** | **H3'-H6 distance** | **PDB structure** | **Nucleotide** | **H2'-H6 distance** | **H3'-H6 distance** |
| 1DDY | U227 | 3 | 3 | 1DDY | U206 | 4 | 2.8 |
| 1DDY | U223 | 2.5 | 4.1 | 5C7W | U69 | 4 | 2.9 |
| 5C7W | U47 | 2 | 4.1 | 5C7W | U67 | 4.1 | 3.4 |
| 5C7W | U48 | 1.9 | 4 | 5DH6 | U8 | 3.7 | 2.9 |
| 5DH6 | U20 | 2.6 | 4.2 | 5DH6 | U24 | 3.9 | 3 |
| 5DH6 | U33 | 2.3 | 4.3 | 5BTP | U68 | 3.8 | 2.6 |
| 5BTP | U31 | 2.6 | 4.4 | 5BTP | U10 | 3.9 | 2.8 |
| 5BTP | U32 | 2.7 | 4.4 | 5BTP | U12 | 4 | 2.9 |
| 4ZNP | U72 | 3.8 | 5.4 | 4ZNP | U41 | 4 | 2.9 |
| 4XNR | U34 | 2.6 | 4.3 | 4ZNP | U43 | 3.9 | 3 |
| 4XNR | U36 | 2.3 | 4.3 | 4ZNP | U66 | 4.2 | 3.1 |
| 1I9V | U7 | 2.6 | 4.3 | 4XNR | U70 | 4 | 3 |
| 1I9V | U17 | 3.5 | 4.9 | 4XNR | U71 | 3.9 | 2.7 |
| 2A43 | U17 | 2.4 | 4.2 | 1I9V | U69 | 4 | 3 |
| 2EES | U22 | 2.6 | 4.4 | 1I9V | U41 | 4.1 | 3.2 |
| 2EES | U36 | 2.4 | 4.4 | 2EES | U25 | 4.2 | 2.9 |
| 2HO6 | U48 | 2.6 | 4.5 | 2EES | U69 | 4 | 2.8 |
| 2HO6 | U79 | 3.1 | 4.4 | 2HO6 | U72 | 3.9 | 2.7 |
| 2GDI | U62 | 2.4 | 4.3 | 2HO6 | U134 | 3.8 | 2.7 |
| 2GDI | U79 | 3 | 3.4 | 2HO6 | U100 | 4.1 | 3.1 |
|  | **AVG:** | 2.645 | 4.265 |  | **AVG:** | 3.975 | 2.92 |
|  | **STDEV:** | 0.455925894 | 0.477135091 |  | **STDEV:** | 0.129269201 | 0.190843005 |
|  | **N:** | 20 | 20 |  | **N:** | 20 | 20 |
|  | **SE:** | 0.101948129 | 0.10669065 |  | **SE:** | 0.028905472 | 0.042673793 |
