## Supplemental Methods for "Structural impact of 3-methylcytosine modification on the anticodon stem of a neuronally-enriched arginine tRNA"

RESP Fitting Procedure:

Step 1:	Create a m3C molecule (Figure S1) using xleap and save it in mol2 format (m3C.mol2). Make sure all bonds are specified under **@<TRIPOS>BOND** section in the mol2 file.

Step 2: Run Amber antechamber to make Gaussian 16 input file. QM optimization is run using Gaussian 16.

**antechamber -fi mol2 -fo gzmat -i m3C.mol2 -o m3C.gau**

**g16 m3C.gau m3C.out**

Step 3: Electrostatic Potential (ESP) around the molecule is extracted using antechamber.

**antechamber -fi gout -fo mol2 -i m3C.out -o m3C_new.mol2 -c resp**

Antechamber does resp fitting in this step and generates files, including a new mol2 file (m3C_new.mol2) with fit charges, ANTECHAMBER.ESP, ANTECHAMBER_RESP1.IN, and ANTECHAMBER_RESP2.IN . This fit does not include the required charge fitting constraints. ANTECHAMBER.ESP, ANTECHAMBER_RESP1.IN, and ANTECHAMBER_RESP2.IN are needed to perform the constrained RESP fit. These are m3C.esp, m3C_step1.respin and m3C_step2.respin in the following text.

Step 4: RESP fitting is a 2-step process. Modify the *respin to include fitting constraints (Figure S1 and S2). The specifics of the *.respin file can be found in the Electrostatic Parameterization with py_resp.py in the AMBER 24 manual. The charge center to fit and its corresponding charge constraints are mentioned at the end of the *respin file:

Fitting Constraints derivation:

Total charge of atoms in cytosine marked in black (Figure S2) = **-0.116**

Total charge of cytosine = **0**

**Total charge of atoms of cytosine marked in red (Figure S2) = +0.116**

**QM structure has charge = +1**

**The total charge of atoms in m3C marked in red (Figure S1) = +1.116**

The RNA OL3 charges for the backbone atoms were listed in m3C_step1.qin. The order of charges needs to match the order of the atoms in the m3C.mol2 file. This requires manual editing of the qin file. For the backbone atoms except C1', partial charges from RNA.OL3 forcefield is listed. These charges are not fit. The remaining charges are kept as dummy values, so that the total sum of all these charges is +1. These remaining charges will be fit according to the fitting constraints mentioned above.

Run the two stages of the resp fitting:

**resp -O -i m3C_step1.respin -o m3C_step1.respout -q m3C_step1.qin -e m3C.ESP -t m3C_step1.crg**

**resp -O -i m3C_step2.respin -o m3C_step2.respout -q m3C_step1.crg -e m3C.ESP -t m3C_step2.crg**

Step 5: As the m3C has flanking residues, to implement the charges, a new mol2 file was created (m3C_P.mol2). The HO5' was replaced with a PO_2_ group and HO3' was removed (Figure S3). Create a new charge list file m3C_fit_P.crg with charges combining RESP fit charges of m3C for base and C1' and RNA.OL3 charges for the remaining backbone atoms. The charges should be in the same order as the atoms in the new m3C_P.mol2 file. Run antechamber, followed by parmchk2 to create a frcmod file. The frcmod file in AMBER is a custom parameter file that defines forcefield modification and additional parameters like bond, angle, dihedral, improper rotation and nonbonded parameters according to RNA.OL3 atom types (Figure S3).

**antechamber -i m3C_P.mol2 -fi mol2 -o m3C_P_charged.mol2 -fo mol2 -c rc -cf m3C_fit_P.crg -at amber**

**parmchk2 -i m3C_P_charged.mol2 -f mol2 -o m3C_fit.frcmod**

Step 6: Using the m3C_P_charged.mol2 and m3C_fit.frcmod file in leap, create a lib file. The lib file provides parameters, structural details, and connectivity information for the m3C residue. It includes the residue name, the atom names in the PDB file that correspond to the atom types, the atomic mass, the partial charges, the connectivity information within the residue and with neighboring residues, and the position of each atom in 3D space.

Use the following commands to invoke leap and then to run the steps in the leap:

**tleap**

**>> loadamberparams m3C_fit.frcmod**

**>> m3C = loadMol2 m3C_P_charged.mol2**

**>> saveOff m3C m3C_fit.lib**

**>> quit**

In the lib file, edit ***!entry.m3C.unit.residueconnect*** and ***!entry.m3C.unit.connect*** to assign the P atom and O3' atom to be the connecting atoms to adjacent residues (Use atom numbers of P and O3' according to the m3C_P_charged.mol2 file). Check if all the required bonds and bond numbers are specified correctly in the ***!entry.m3C.unit.connectivity*** section.

Step 7: Combine the frcmod file and lib file to a leaprc.m3C.

**echo "m3C = loadAmberParams m3C_fit.frcmod" > leaprc.m3C**

**echo "loadOFF m3C_fit.lib" >> leaprc.m3C**

Step 8: Load this leaprc file along with forcefield to use these parameters in simulations/annealing in leap:

**tleap**

**>> source leaprc.RNA.OL3**

**>> source leaprm.m3C**

**>> molecule = loadpdb m3C_ArgAC.pdb**

**>> saveamberparm molecule m3C_ArgAC.prmtop m3C_ArgAC.rst7**

**>> quit**

Figures:

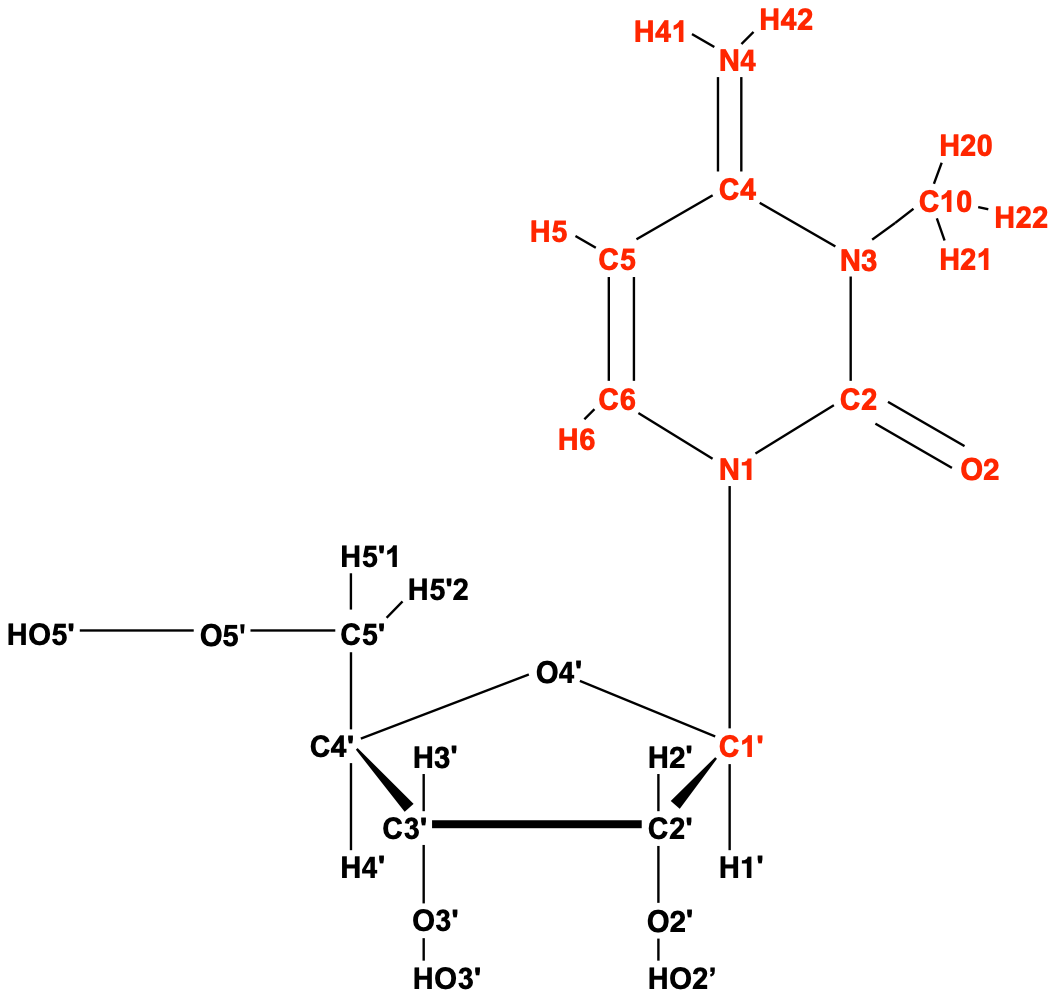

Figure S1: Structure of m3C (m3C.mol2) used to run QM calculation with a total charge of +1.

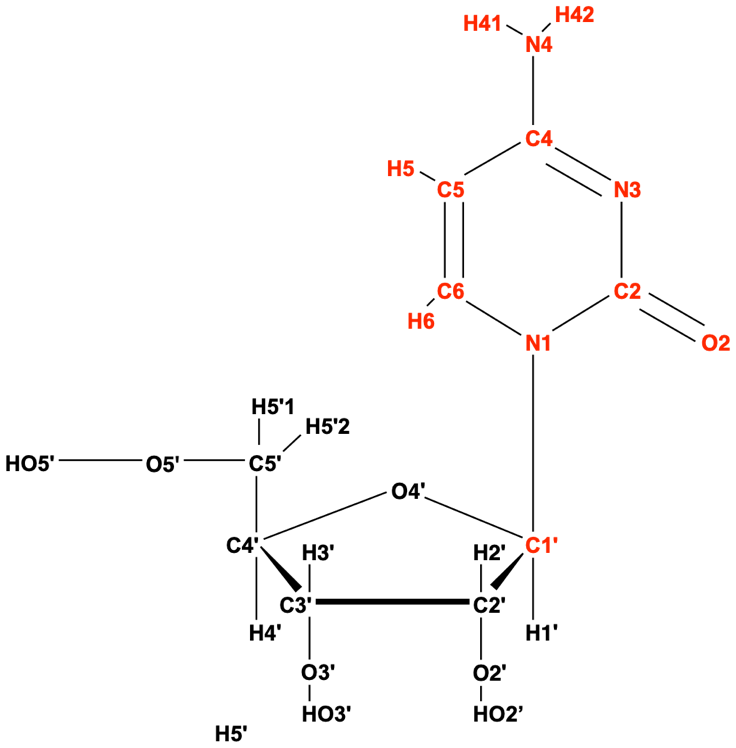

Figure S2: Charges of atoms marked in red (in Figure S1) were fit and the remaining were equated to cytosine RNA OL3 forcefield charges (atoms marked in black).

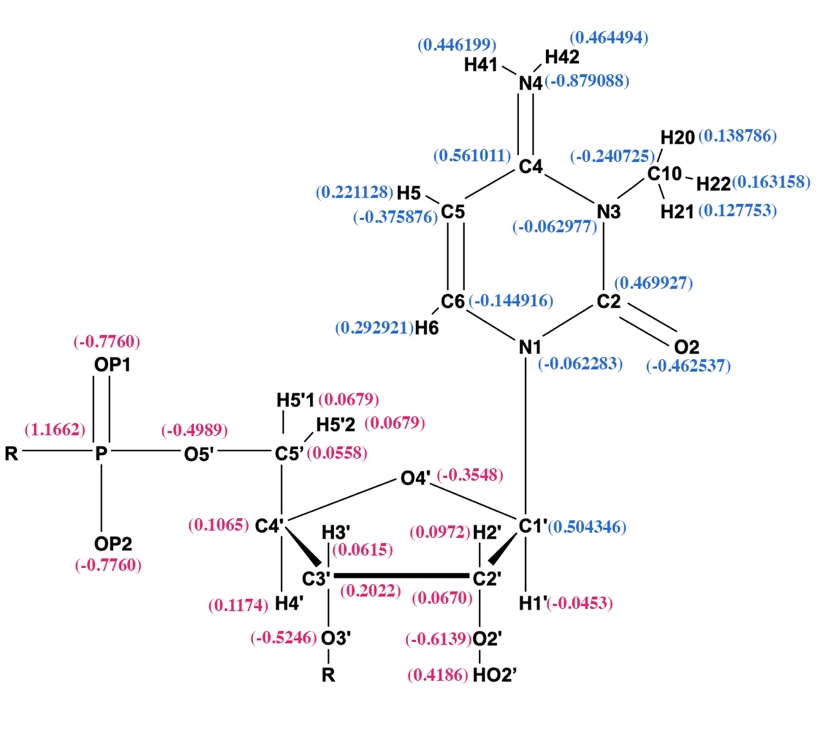

Figure S3: Structure of m3C_P.mol2 used in the annealing step. The RESP fit charges are marked in blue and the charges assigned from RNA.OL3 are marked as purple.
